# Three-dimensional mapping identifies distinct vascular niches for myelopoiesis

**DOI:** 10.1101/2020.04.02.014548

**Authors:** Jizhou Zhang, Qingqing Wu, Courtney B. Johnson, Andre Olsson, Anastasiya Slaughter, Margot May, Benjamin Weinhaus, Angelo D’Alessandro, James Douglas Engel, Jean X. Jiang, J. Matthew Koffron, L. Frank Huang, Nathan Salomonis, H. Leighton Grimes, Daniel Lucas

## Abstract

In contrast to virtually all other tissues in the body the anatomy of differentiation in the bone marrow remains unknown. This is due to the lack of strategies to examine blood cell production in situ, which are required to better understand differentiation, lineage commitment decisions, and to define how spatial organizing cues inform tissue function. Here we developed imaging approaches to map all myeloid cells in whole bones and generated 3D atlases of granulocyte and monocyte/dendritic cell differentiation during homeostasis. We found that myeloid progenitors leave the hematopoietic stem cell niche during differentiation. Granulocyte and monocyte dendritic cell progenitors (MDP) do not interact, instead they localize to different sinusoids where they give rise to clusters of immature cells. MDP cluster with Ly6C^lo^ monocytes and conventional dendritic cells; these localize to a unique subset of colony stimulating factor 1 (CSF1, the major regulator of monopoiesis^1^) -expressing sinusoids. Csf1 deletion in the vasculature disrupted the MDP clusters and their interaction with sinusoids, leading to reduced MDP numbers and differentiation ability, with subsequent loss of peripheral Ly6C^lo^ monocytes and dendritic cells. These data indicate that there is a specific spatial organization of definitive hematopoiesis and that local cues produced by distinct blood vessels are responsible for this organization. These maps provide a blueprint for in situ analyses of hematopoiesis in blood disorders.

## Introduction

In the bone marrow (BM) hematopoietic stem cells (HSC) and multipotent progenitors progressively differentiate and commit to give rise to lineage-specific progenitors that then generate all major blood cell lineages. Hematopoietic differentiation has been studied in great detail using multiple approaches including scRNAseq^2-4^, and in vivo lineage tracing/barcoding^5-10^. Even though these approaches provide critical insight on the regulation and pathways of differentiation they require destruction of the organization of the originating cells in tissue. Most of our knowledge of the anatomy of blood differentiation still derives from classical studies using light or electron microscopy that relied on nuclear and cellular morphology to identify the differentiating cells^11-14^. These studies suggested regional organization of differentiation. A classic example was the discovery that erythropoiesis is organized around macrophage islands near sinusoids^12^. Because the vast majority of BM cells cannot be uniquely identified using these criteria the functional anatomy of differentiation and the structures that regulate specific cell types remain poorly understood.

Confocal imaging analyses of HSC and other progenitors have been invaluable in identifying the cellular components and organization of the niches that regulate them^15-18^. The HSC niche is a multicellular structure composed of blood vessels, perivascular cells (identified using *Cxcl12, LepR* and/or *Nestin* reporter mice), megakaryocytes, and other cells^19,20^. Common lymphoid progenitors (CLP) and erythroid progenitors also localize to-and are regulated by- the CXCL12-producing perivascular cells that support HSC^15,17,21,22^ suggesting that they share the same niche. However, colocalization of HSC and committed progenitors has not yet been demonstrated and some common lymphoid progenitors localize to the endosteum and are regulated by osteoblastic cells^21,23^. These suggest that some progenitors leave the HSC niche. Whether lineage committed progenitors share the same niche as HSC or are regulated by distinct niches remains an open question.

Here we report the mapping of myelopoiesis in situ; we demonstrate that myeloid progenitors abandon the HSC niche upon differentiation, and we identify a unique subset of sinusoids that provide a niche for the Ly6C^lo^ monocyte and dendritic cell production.

## Results

### Development of imaging approaches to analyze myelopoiesis in situ

Classically, most hematopoietic progenitors have been identified using combinations of cell surface markers that allowed their purification via fluorescence activated cell sorting (FACS). Although not surprisingly scRNAseq showed that FACS-purified progenitors are heterogeneous^2-4^, cell surface markers remain invaluable to identify populations highly enriched in specific progenitors with unique developmental potential^2,24^.

In myelopoiesis a common myeloid progenitor (CMP) can differentiate into monocyte dendritic cell progenitors (MDP) or granulocyte-monocyte progenitors (GMP). MDP can differentiate into common monocyte progenitors (cMoP) or common dendritic progenitors that give rise to monocytes or dendritic cells (DC) respectively (Fig. 1a). GMP can also generate neutrophil-like monocytes via an intermediate monocyte progenitor (MP) and neutrophils via a granulocyte progenitor (GP) that gives rise to pre-, immature, and mature neutrophils^24,25^ (Fig. 1a). cMoP and MP express the same cell surface markers and can only be distinguished transcriptionally^24^. For simplicity, we will refer here to the population containing both cMoP and MP as MOP.

**Figure 1.**
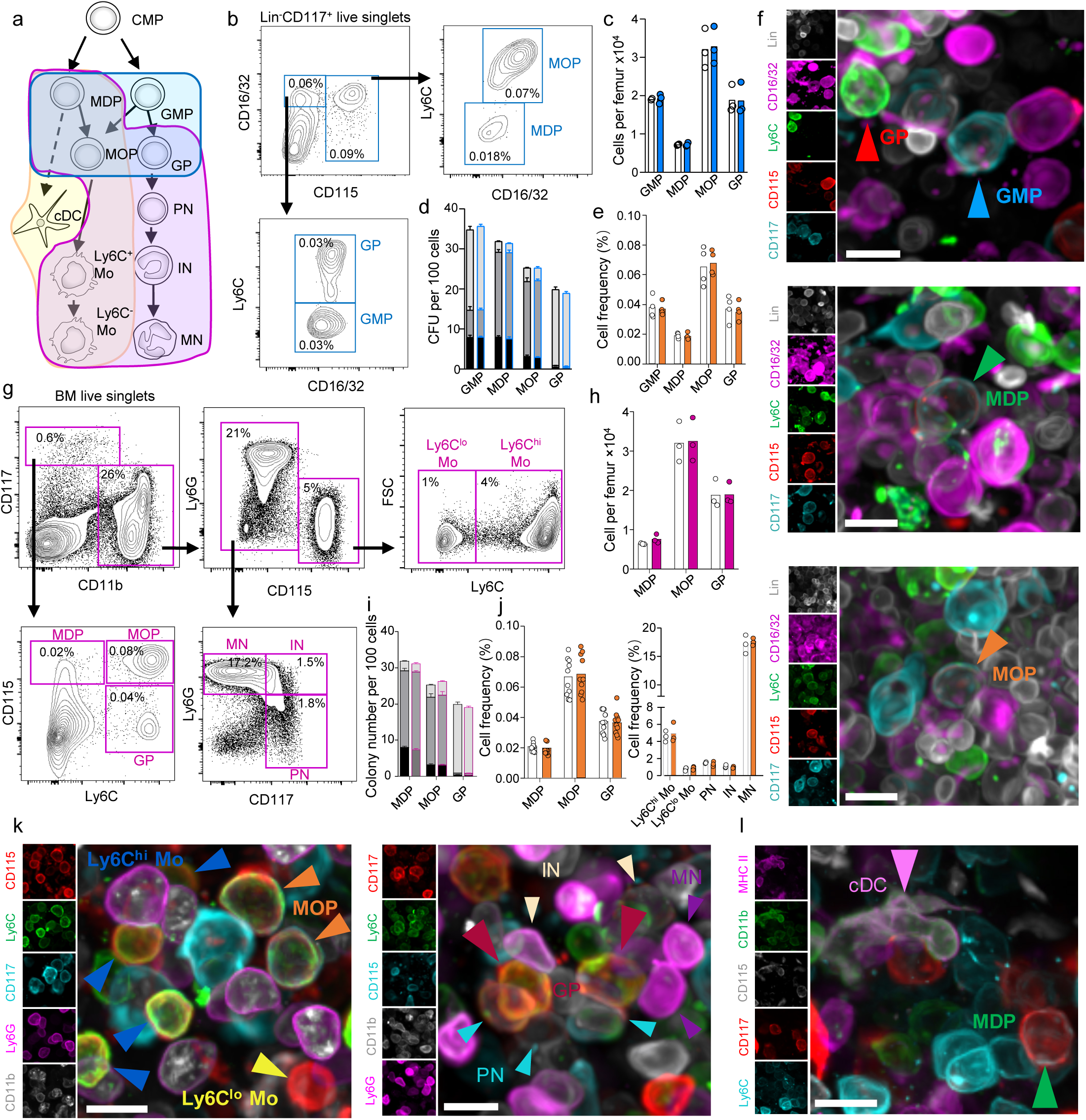
**a**. Schematic representation of myelopoiesis. The three shaded areas indicate the three different stains used to image the different populations. CMP: Common myeloid progenitor. MDP: Monocyte dendritic cell progenitor. GMP: granulocyte monocyte progenitor. MOP: gate containing cMoP and MP progenitors. GP: granulocyte progenitor. cDC: conventional dendritic cell. Mo: monocyte. PN: Pre-Neutrophil. IN: Immature Neutrophil. MN: Mature neutrophil. **b**. FACS gating strategy for isolation and imaging of the indicated progenitors. The Lineage panel contains antibodies against Ly6G, CD11b, Ter119, B220 and CD3. **c, d**. Frequency (c) and colony forming activity (d, black: CFU-GM, grey: CFU-M, white: CFU-G; n = 3 mice) of the indicated progenitors using previously described strategies^24,26^ (white bar or black border) or the one described in b (blue bar or blue border). **e**. Frequency of total BM cells for each of the indicated populations when detected by FACS (white) or imaging (orange). **f**. Representative images showing the different progenitors in the BM. **g**. FACS gating strategy for isolation and imaging of the indicated cells. **h, i**. Frequency (h) and colony forming activity (i, black: CFU-GM, grey: CFU-M, white: CFU-G; n = 3 mice) of the indicated progenitors using previously described strategies^24,26^ (white bar/black border) or the one described in g (red bar/red border). **j**. Frequency of total BM cells for each of the indicated populations when detected by FACS (white) or imaging (orange). **k, l**. Representative images showing detection of mature myeloid cells in the bone marrow. For all images the Scale bar = 10µm. For all histograms one dot corresponds to one mouse.

To examine myelopoiesis in situ we developed three antibody stains (Figure1a shaded areas and Supplementary table 1). For the first stain (Fig. 1a blue area) we used CD117, CD115 and Ly6C to identify MDP and MOP as described previously (^26^ and Extended Data Fig. 1a). These three markers can be combined with a Lineage panel and CD16/32 to simultaneously detect MDP, GMP, MOP and GP by FACS^24^ and imaging (Fig. 1a-f and Extended Data Fig. 1b). We found the same frequencies and colony-forming activities (Fig. 1c, d) between the cells in these gates and the established FACS stains^24,26^ shown in Extended Data Fig. 1a, b indicating that they label the same populations. We found identical frequencies when using imaging or FACS (Fig. 1e, f), demonstrating that imaging detected that entire population of progenitors in the BM. Ly6C staining additionally allowed visualization of arterioles in the BM (Extended Data Fig. 1c).

For the second stain we realized that it was possible to replace the Lineage panel with CD11b to identify MDP, MOP and GP (Fig. 1a purple area, Fig. 1g and Extended Data Fig. 1d, e) by flow cytometry. These gates yielded identical frequencies (Fig. 1h) and colony forming activity (Fig. 1i) as shown previously (^24,26^ and Extended Fig. 1a, b). Other studies used *Cx3cr1-gfp* reporter mice to identify MDP and MOP^27^. To further validate this novel approach, we confirmed that MDP and MOP using this strategy were uniformly GFP^+^ in *Cx3cr1-gfp* (Extended Data Fig. 1f). GP and MOP can be distinguished based on high *Gfi1* and *Irf8* expression respectively^2,24^ and we confirmed differential expression of these factors in GP and MOP by qRT-PCR and using *Gfi1* reporter mice (Extended Data Fig. 1g, h). CD11b staining also allowed simultaneous detection of Ly6C^hi^ and Ly6C^lo^ monocytes (Fig. 1g). We replaced CD16/32 with Ly6G antibodies allowing identification of CD11b^+^CD117^dim^CD115^-^Ly6G^lo^ pre-neutrophils^25^ (enriched in metamyelocytes and very immature band cells); CD11b^+^CD117^dim^CD115^-^Ly6G^hi^ immature neutrophils (enriched in banded neutrophils); and CD117^-^Ly6G^hi^ mature neutrophils^25^ (Fig. 1g and Extended Data Fig. 1i). In the imaging analyses we were able to simultaneously detect these 8 populations (Fig. 1a, purple area) at comparable frequencies to those detected by FACS (Fig. 1j, k).

Dendritic cells can be imaged as reticulated Cx3cr1*-*GFP^+^ or Cx3cr1*-*GFP^+^MHCII^+^ cells in *Cx3cr1-gfp* reporter mice^28,29^. All reticulated GFP^+^ cells were also MHCII^+^ indicating that MHCII and cell shape are sufficient to unambiguously identify DC (Extended Data Fig. 1j). Reticulated MHCII^+^ cells are conventional dendritic cells (cDC) as they are positive for CD11b and negative for B220 and CD8 (Extended Data Fig. 1k). For the third stain we replaced Ly6G with MHCII in the stain in Fig. 1g to simultaneously detect MDP, MOP, Ly6C^hi^ and Ly6C^lo^ monocytes, and cDC in the BM (Fig. 1a yellow area, Fig. 1l, and data not shown).

### 3D atlases of steady state myelopoiesis

We used the stains above to manually map the 3D position of all myeloid lineage cells in the sternum (>60,000 cells analyzed in 38 sterna) and assess the relationships between progenitors and their offspring with single cell resolution in situ. Due to the complexity of the stains we replaced each different cell with a color-coded sphere centered in the cell to better visualize differentiation (Fig. 2a and Extended Data Fig. 2a and Supplementary Video 1). We also obtained the X, Y, and Z coordinates for every cell, and then used these data to quantify the distance to all other cells. To test whether the spatial relationships observed were specific we compared them to those predicted from a random distribution. For this we obtained the X, Y and Z coordinates for every hematopoietic cell in the sternum (detected using αCD45 and αTer119 Extended Data Fig. 2a). We then used these coordinates and randomizing software to randomly place each cell type, at the exact same frequencies found in vivo, throughout the BM. Each random distribution simulation was repeated between 100-200 times.

**Figure 2.**
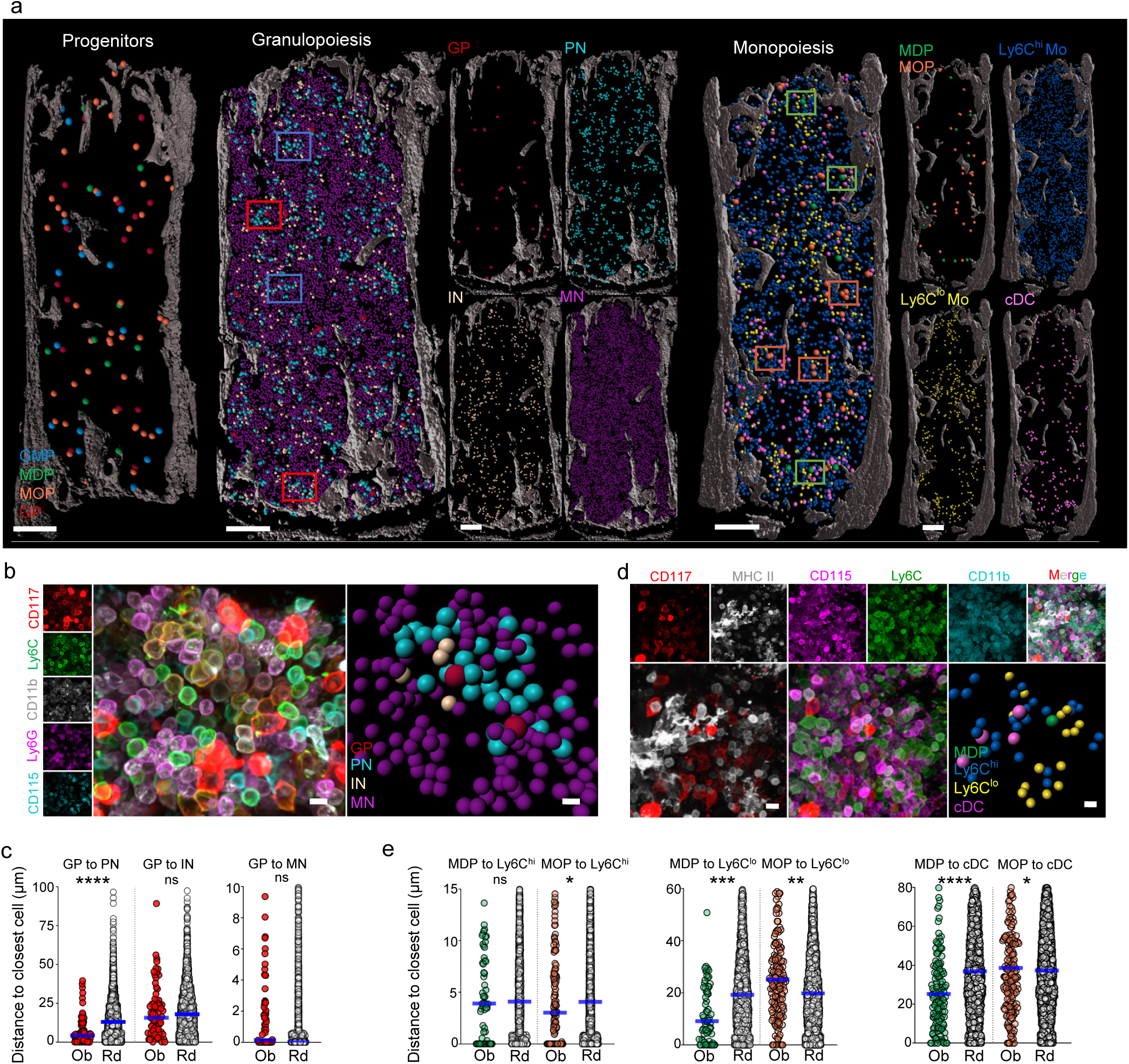
**a**. Maps showing the location of the indicated cells in the bone marrow. Each dot corresponds to one cell. The radius of each dot is 3x (for progenitors) 1.5x (Mature Neutrophils) or 2x (all other cells) the average radius of the replaced cell. Squares highlight representative clusters around GP (red squares) or clusters without GP (blue squares). Green squares highlight areas around MDP and orange squares highlight clusters of 2-3 MOP. Scale bar = 200μm. **b**. Representative image showing neutrophil differentiation around a GP. Each dot radius corresponds to the average radius of the replaced cell. Scale bar = 10μm. **c**. Histograms showing the observed (red) and random (white) distribution of distances from each GP to the closest indicated cell (n = 75 GP in 3 sterna of 3 mice). **d**. Representative image showing Ly6C^lo^ monocyte and cDC localization near MDP. Scale bar = 10μm. Each dot radius corresponds to the average radius of the replaced cell. **e**. Histograms showing the distance from each MDP (green dots) or MOP (orange dots) to the closest indicated cell (MDP-Ly6C^hi^ monocyte, n = 67 MDP from 6 sterna of 4 mice; MOP-Ly6C^hi^ monocyte, n = 137 MOP from 3 sterna of 3 mice; MDP-Ly6C^lo^ monocyte, n = 67 MDP from 6 sterna of 4 mice; MOP-Ly6C^lo^ monocyte, n = 171 MOP from 4 sterna of 4 mice; MDP-cDC, n = 139 MDP from 11 sterna of 6 mice; MOP-cDC, n = 200 MOP from 5 sterna of 4 mice). In all graphs one dot corresponds to one cell. Horizontal blue bars indicate the median distance.

Despite their proximity in the myeloid differentiation tree (Fig. 1a and ^24^) GMP, MDP, MOP and GP are interspersed throughout the marrow and appear to minimally interact (Fig. 2a and Extended Data Fig. 2b and Supplementary Video 2). This is in agreement with the data from a previous study that showed that Lin^-^CD117^+^CD16/32^+^ progenitors (which contain GMP, GP and MOP) were interspersed through the central BM^16^. Our data shows GP are closer to other GP than predicted from random distribution (Fig. 2a, b and Extended Data Fig. 2b) and 1 or 2 GP generate pre-neutrophils that are tightly (median distance=4.16 μm) clustered near the GP (Fig. 2a, red squares and Fig. 2b, c and Supplementary Videos 3, 4). Immature neutrophils are found in the periphery of the cluster and migrate away as they differentiate into mature neutrophils which are evenly distributed throughout the BM and do not specifically associate with GP (Fig. 2a-c). Numerous clusters of pre-neutrophils and immature neutrophils do not contain GP. The cells in these clusters are farther away from each other than in the clusters centered on GP (Fig. 2a blue squares and Extended Data Fig. 2c) suggesting that when GP differentiate, the cluster disaggregates.

The next analyses showed that MOP are significantly closer to each other than predicted randomly, and up to 3 MOP can be detected nearby one another (Fig. 2a orange squares and Extended Data Fig. 2b and Supplementary Video 5). MDP are also closer to each other than predicted by random chance but are not close enough to be adjacent (Fig. 2a green squares and Extended Data Fig. 2b and Supplementary Video 5). Ly6C^hi^ monocytes are spread through the central BM (Fig. 2a) and are minimally enriched near MDP and MOP (Fig. 2a, d, e), suggesting that Ly6C^hi^ monocytes quickly move away from their progenitors after differentiation. In contrast Ly6C^lo^ monocytes form loose clusters ((median distance=12.26 μm; Fig. 2a and Extended Data Fig. 2d) that are selectively enriched near MDP but depleted near MOP (Fig. 2a, d, e and Extended Data Fig. 2e, f and Supplementary Video 6). In agreement with previous studies^28,29^ we found that cDC form clusters (Fig. 2a and Extended Data Fig. 2g). These clusters are enriched near MDP but not MOP (Fig. 2e and Extended Data Fig. 2h, I and Supplementary Video 6). These indicate regional compartmentalization of Ly6C^lo^ monocyte and cDC production around MDP.

### Granulopoiesis and monopoiesis segregate to different sinusoidal locations far away from HSC

GP identify areas of granulopoiesis (Fig. 2a-c) and MDP identify areas of Ly6C^lo^ monocyte and cDC production (Fig. 2a, d, e) respectively, but these progenitors do not colocalize (Fig. 2a and Extended Data Fig. 2b) suggesting that generation of neutrophils, monocytes and dendritic cells does not occur in the same location. In the experiments shown in Fig. 1g we noticed that pre-neutrophils and immature neutrophils are the only CD117^+^ cells in the CD11b^+^CD115^-^ gate (Fig. 1g). This allowed us to simultaneously image a population containing both cell types together with Ly6C^lo^ monocytes and dendritic cells (Extended Data Fig. 3a). We found that Ly6C^lo^ monocyte and cDC are farther away from pre- and immature neutrophils than predicted from random distributions (Fig. 3a, b), demonstrating that granulopoiesis and Ly6C^lo^ mono- and DC-poiesis take place in different bone marrow locations.

**Figure 3.**
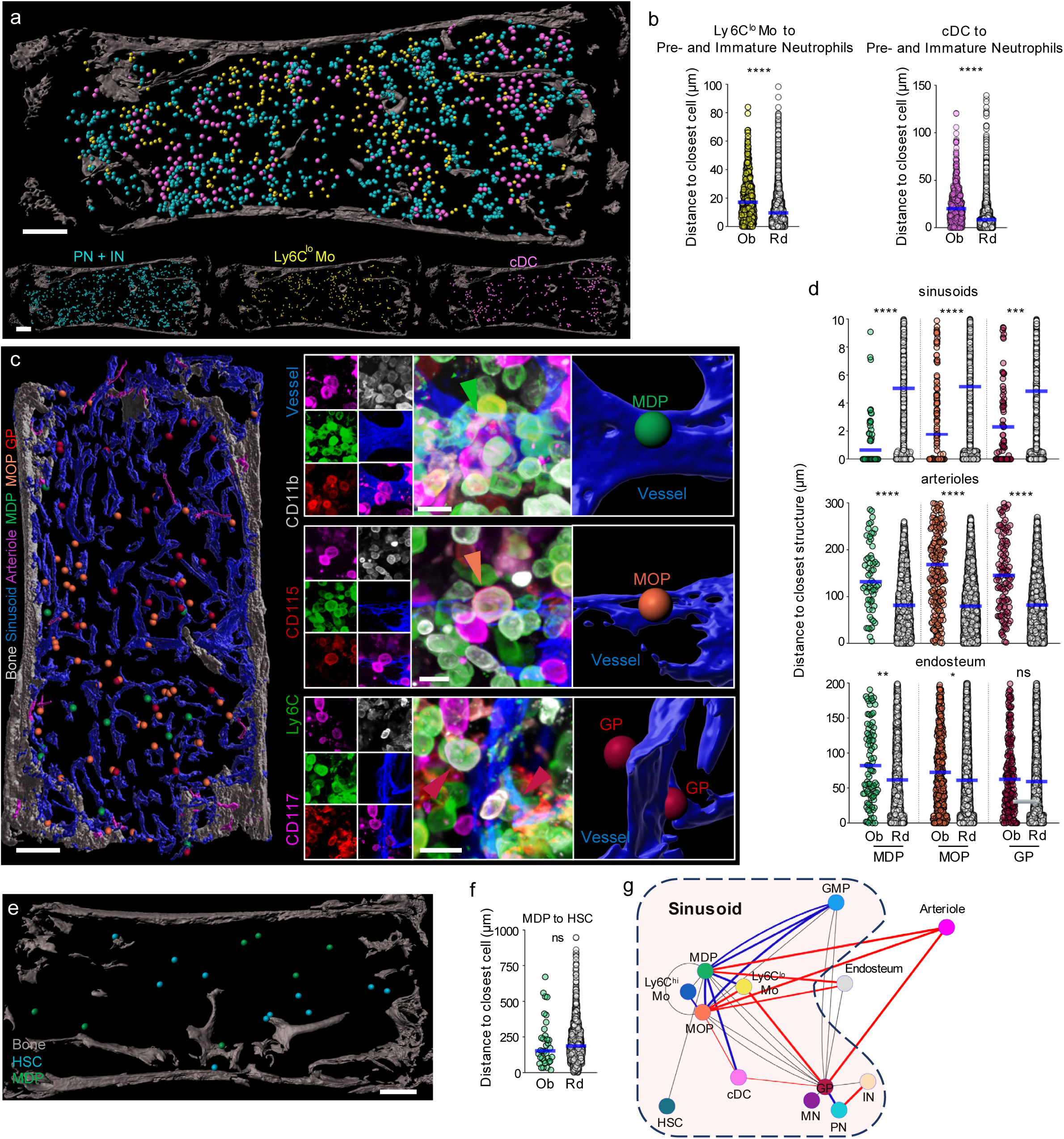
**a**. Map showing the location of the indicated cells in the bone marrow. Each dot corresponds to one cell. Note that the radius of each dot is 2x the average radius of the cell. Scale bar = 200μm. **b**. Histograms showing the distance from each Ly6C^lo^ Mo (yellow dots) or cDC (pink dots) and their random simulation (white dots) to the closest Pre- or immature neutrophil (n = 500 Ly6C^lo^ Mo and n = 727 cDC, from 3 sterna of 3 mice). **c**. Map (left panel) and high-power images (right panels) showing the interactions between MDP, MOP and GP with the bone (white), sinusoids (dark blue) or arterioles (pink). Each dot corresponds to one cell, for the map the dot radius is 3x whereas for the right panels the dot radius is the same as the average radius of the replaced cell. Scale bars are 200μm and 10μm. **d**. Histograms showing the distance from each MDP (green dots), MOP (orange dots), GP (red dots) or random distribution (white dots) to the closest indicated structure (distances to sinusoid and arterioles, n= 62 MDP from 6 sterna of 6 mice; n = 218 MOP, and n = 114 GP from 5 sterna of 5 mice; Distance to bone, n = 98 MDP, n = 410 MOP, n = 217 GP, from 9 sterna of 6 mice). **e**. Map showing the location of HSC and MDP in the bone marrow. Each dot corresponds to one cell and the dot radius is 3x the average cell radius. **f**. Histograms showing the distance from each MDP (green dots) or the random simulation (white dots) to the closest HSC (n = 34 MDP from 4 sterna of 3 mice). **g**. Graph summarizing the distances between the different cell populations examined in this study. Each dot corresponds to one type of cell. We connect cells with lines that indicate if they are significantly closer (blue line) or farther away (red line) or no specific interaction (grey line) when compared to random distributions. If no line is drawn, then we did not examine the interactions between these two cell types. Thicker lines indicate lower p values when testing for the significance of the interaction.

Most HSC localize to sinusoids, whereas smaller fractions associate with arterioles and the endosteal surface^30-33^. These structures provide different signals such as CXCL12, SCF, and IL7 that maintain and regulate HSC^19,20^, CLP and erythroid progenitors^15,17,21-23^. We found that MDP, MOP and GP are farther away from arterioles, show no specific interaction with the endosteum, and are closer to sinusoids when compared to random simulations (Fig. 3c, d and Supplementary Videos 7-9). These data indicate that sinusoids are the site of myelopoiesis. Since granulopoiesis and mono/DCpoiesis do not overlap (Fig. 3a, b and Extended Data Fig. 2b), these results raised the possibility that different sinusoids produce unique signals to regulate specific myeloid progenitors.

MDP can be defined as Lin^-^CD117^+^CD115^+^Ly6C^-^ cells whereas HSC are characterized as Lin^-^ CD117^+^CD48^-^CD41^dim^CD150^+^ cells^18^, so we combined these markers to simultaneously image the cells (Extended Data Fig. 3b). GP and MOP are the only Ly6C^+^ cells in the Lin^-^CD117^+^ gate (Extended Data Fig. 1e) allowing imaging with HSC (Extended Data Fig. 3c). We found no spatial relationship between MDP and HSC (Fig. 3e, f) or the mixed population containing GP and MOP (Extended Data Fig. 3c-e) when compared to random distributions. These results indicate that MDP, GP and MOP - or their upstream progenitors - abandon the HSC niche during differentiation. Fig. 3g provides a summary of the distances and specific interactions between the different cells and structures analyzed in this study.

### A distinct subset of CSF1^+^ sinusoids regulates MDP, Ly6C^lo^ monocytes and cDC to organize myelopoiesis

Colony stimulating factor 1 (CSF1, also known as macrophage colony stimulating factor, M-CSF) is produced by non-hematopoietic cells and is necessary for monocyte, macrophage and osteoclast generation. We reclustered and analyzed published BM scRNA-Seq datasets^34^ to assess *Csf1* expression. Almost all *LepR*-expressing perivascular cells and a small subset of endothelial cells expressed *Csf1* (Fig. 4a and Extended Data Fig. 4a). We bred *Csf1*^*fl/-*^ mice^35^ with *LepR-cre*^*22,36*^ or *Cdh5-cre*^*37*^ mice to conditionally inactivate the *Csf1* gene in perivascular or endothelial cells, respectively (Fig. 4b). Despite efficient deletion (Extended Data Fig. 4b), conditional *Csf1* ablation in LepR^+^ perivascular cells does not impact MDP numbers or function (nor affect any other hematopoietic cell) in bone marrow or blood (Fig. 4c and Extended Data Fig. 4c-f). In contrast *Cdh5-cre:Csf1*^*fl/-*^ (for simplicity *Csf1*^*ΔEC*^) mice showed a significant reduction in MDP numbers that is further compounded by reductions in MDP-derived CFU-M, but not CFU-G or CFU-GM, colonies when compared to control mice (Fig. 4c, d and Extended Data Fig. 4g). GMP and MOP numbers in *Csf1*^*ΔEC*^ mice are identical to those found in controls and their CFU-M activity is minimally impaired (Extended Data Fig. 4g-i). Ly6C^lo^ monocytes and cDC form clusters around MDP (Fig. 2a, d, e). Both cell types show a ∼2-fold reduction in numbers in *Csf1*^*ΔEC*^ mice whereas Ly6C^hi^ monocytes - which do not associate with MDP - are unaffected (Fig. 4e). The reduction in Ly6C^lo^ monocytes persists in peripheral blood (Fig. 4f). *Csf1*^*ΔEC*^ mice do not exhibit changes in BM cellularity, HSC, multipotent progenitors, CMP, GP, or mature cell numbers in bone marrow or blood (Extended Data Fig. 4j-l). HSC express low levels of CSF1R and CSF1 instructs HSC commitment towards myeloid fates^40^. To rigorously assess the impact of endothelial-derived CSF1 on HSC function we performed competitive bone marrow transplants. These experimental data demonstrated no differences in short-term or long-term engraftment or lineage biases between recipients transplanted with *Csf1*^*ΔEC*^ or control bone marrow, indicating that endothelial-derived CSF1 does not regulate HSC function during homeostasis (Extended Data Fig. 4m).

**Figure 4.**
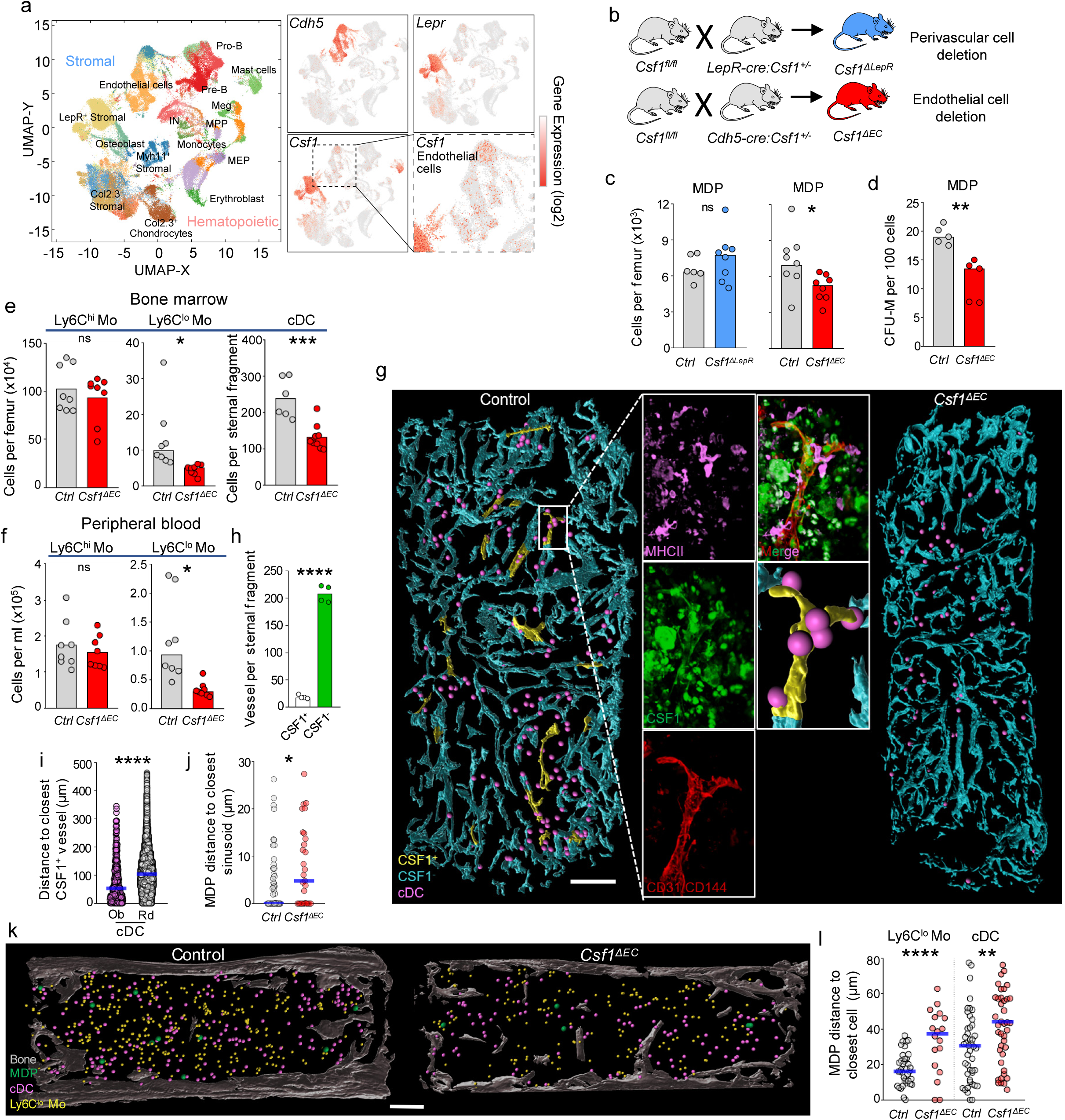
**a**. Left panel: Uniform Manifold Approximation and Projection (UMAP) of bone marrow stromal cells with clusters color-coded. Clusters are reanalyses^42^ of a published dataset^34^. Right panels: *Cdh5, Lepr*, and *Csf1* expression in the bone marrow cells. IN: Immature Neutrophils; MEP: megakaryocyte-erythrocyte progenitors. **b**. Scheme showing the generation of *Csf1*^*ΔLepR*^, *Csf1*^*ΔEC*^ or control (pool of *Cre:Csf1*^*+/-*^, *Csf1*^*+/-*^, and *Csf1*^*fl/*-^) littermates to conditionally delete *Csf1* in perivascular or endothelial cells respectively. **c**. Number of MDP per femur in control, *Csf1*^*ΔLepR*^ or *Csf1*^*ΔEC*^ mice. **d**. Number of CFU-M produced by MDP from control or *Csf1*^*ΔEC*^ mice. **e, f**. Number of Ly6C^hi^ and Ly6C^lo^ monocytes or cDC in the BM (e) or blood (f) of control or *Csf1*^*ΔEC*^ mice. **g**. Map and high-power image showing the location of CSF1^+^ and CSF1^-^ vessels and cDC (pink dots) in wild-type mice. The radius of the dot is 2x the average cDC radius. Scale bar = 200μm. **h**. Number of CSF1^+^ and CSF1^-^ vessels per sternum in wild-type mice. Each dot corresponds to one mouse. **i**. Histograms showing the distance from each cDC (pink dots) or random cell (white dots) to the closest CSF1^+^ vessel (n = 442 cDC from 4 sterna of 3 mice). **j**. Histograms showing the distance from each MDP to the closest vessel in control or *Csf1*^*ΔEC*^ mice (n = 44 MDP from 5 sterna of 3 control mice; n = 29 MDP from 5 sterna of 3 *Csf1*^*ΔEC*^ mice). **k**. Map showing the distribution of MDP, Ly6C^lo^ Mo and cDC in the sternum of control or *Csf1*^*ΔEC*^ mice. Scale bar = 200μm. The radius of the dot is 3x (MDP) or 2x (all other cells) the average radius of the replaced cell. **l**. Histograms showing the distance from each MDP in control (grey dots) or *Csf1*^*ΔEC*^ mice (red dots) to the closest Ly6C^lo^ monocyte or cDC. For MDP-Ly6C^lo^ monocyte, n = 37 MDP from 4 sterna of 3 control mice, n = 18 MDP from 4 sterna of 3 *Csf1*^*ΔEC*^ mice. For MDP-cDC, n = 47 MDP from 6 sterna of 3 control mice, n = 47 MDP from 9 sterna of 3 *Csf1*^*ΔEC*^ mice).

The scRNA-seq analyses suggested that only a small fraction of BM endothelial cells produce *Csf1* (Fig. 4a). Using a polyclonal antibody against CSF1 followed by a HRP-conjugated secondary antibody and tyramide signal amplification, we were able to visualize the rare CSF1^+^ cells that represented 8% of all BM blood vessels (Fig. 4g, h). This labeling was specific as it was almost completely abolished in the *Csf1*^*ΔEC*^ mice (Fig. 4g). The stain was compatible with MHCII antibody allowing detection of cDC (Fig. 4g and Supplementary Video 10) but no other populations as the amplification protocol caused loss of most cell surface markers. cDC are specifically enriched around the CSF1^+^ vessels (Fig. 4g, i and Extended Data Fig. 5a). *Csf1* deletion in the vasculature caused MDP-but not cDC to move away from sinusoids (Fig. 4j and Extended Data Fig. 5b, c); and disrupted the clusters of Ly6C^lo^ monocytes and cDC around MDP (Fig. 4k, l and Extended Data Fig. 5d). Taken together, these data indicate that CSF1^+^ sinusoids are a niche for MDP that regulates Ly6C^lo^ monocytes and cDC production.

## Discussion

The anatomy of differentiation in the bone marrow remains largely unknown. Here we described imaging approaches to map myelopoiesis in situ. Myeloid progenitors do not colocalize with HSC, indicating that they migrate away from HSC niches. Migration seems to be intrinsic to myelopoiesis as – despite sharing a common progenitor - GMP, MDP, MOP and GP are not close to one another. MDP and GP are recruited to different sinusoids where they divide to form clusters containing large numbers of immature cells that leave the cluster as they differentiate. Previous studies showed that 99% of all hematopoietic cells were within 30μm of sinusoids^38^ and that CXCL12 and SCF were widely distributed through these sinusoids^21,22^. These studies raised the possibility that sinusoids are a widespread niche that provides broad support for all types of neighboring cells. The present data indicates that not all sinusoids are equivalent, and that a small subset (8%) of CSF1^+^ sinusoids provide a unique niche responsible for organizing and regulating MDP, monocytes and dendritic cells. The fact that other progenitors (GP, MOP) localize to different sinusoids suggests the existence of additional local niches that will support their differentiation. The data also provides novel insights into how the bone marrow controls progenitor differentiation. CSF1 instructs lineage choice in HSC and GMP^39,40^, is required for MDP and cMoP differentiation into monocytes, drives a fraction of monocytes towards macrophage and osteoclast fates and maintains subsets of monocytes and dendritic cells (reviewed in ^1^). Mice lacking CSF1 or its receptor show an almost complete absence of monocytes and macrophages and the bone marrow cavity is reduced due to depletion of osteoclasts ^35,41^. *Csf1* deletion in the vasculature yields a much milder phenotype as it impacts MDP, Ly6C^lo^ monocytes and cDC but no other hematopoietic cell type. This indicates that other CSF1 sources function in hematopoiesis and suggest that the bone marrow controls production of specific lineages by compartmentalizing cytokine production to unique locations.

## KEY SOURCE TABLE

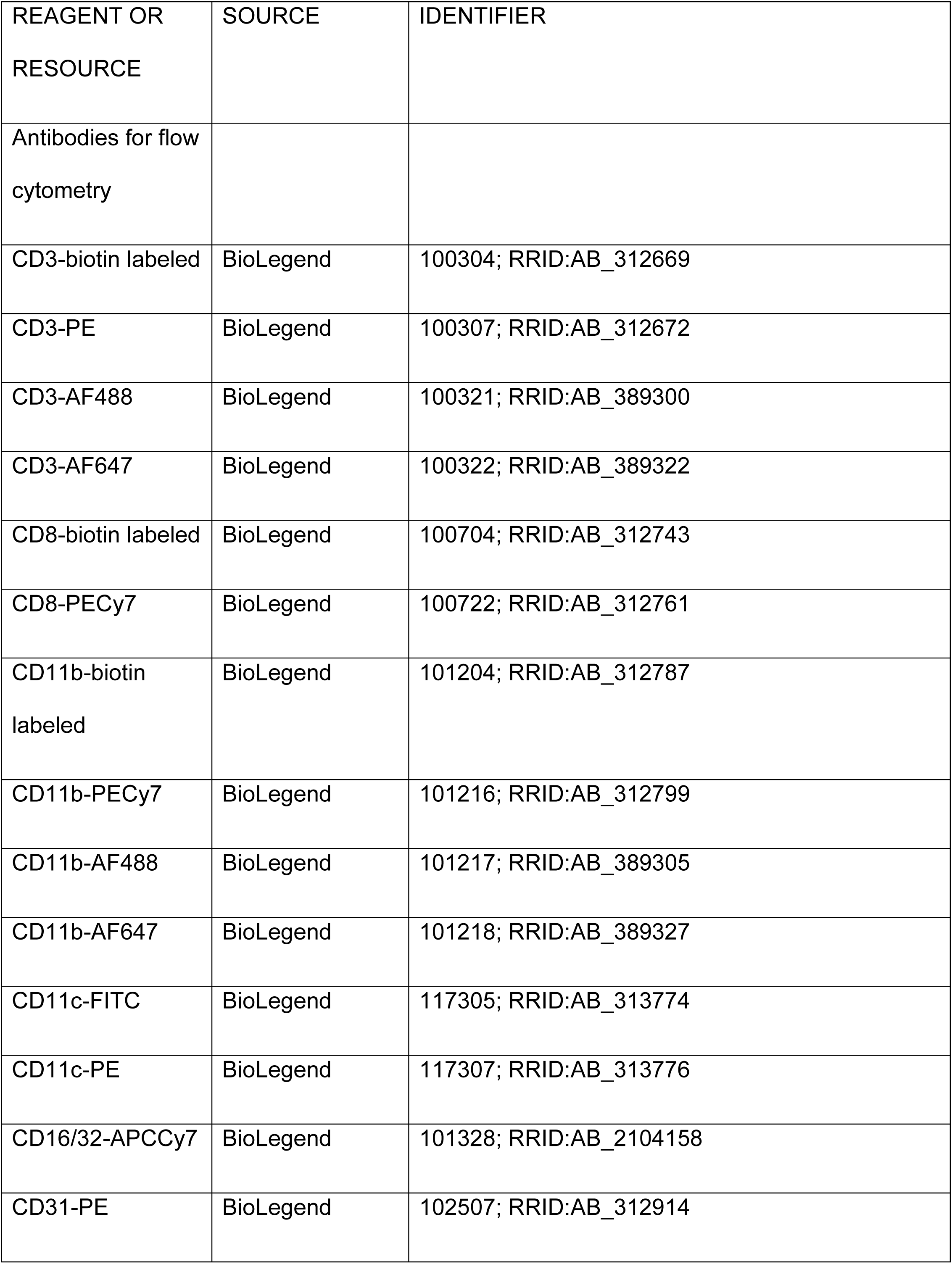
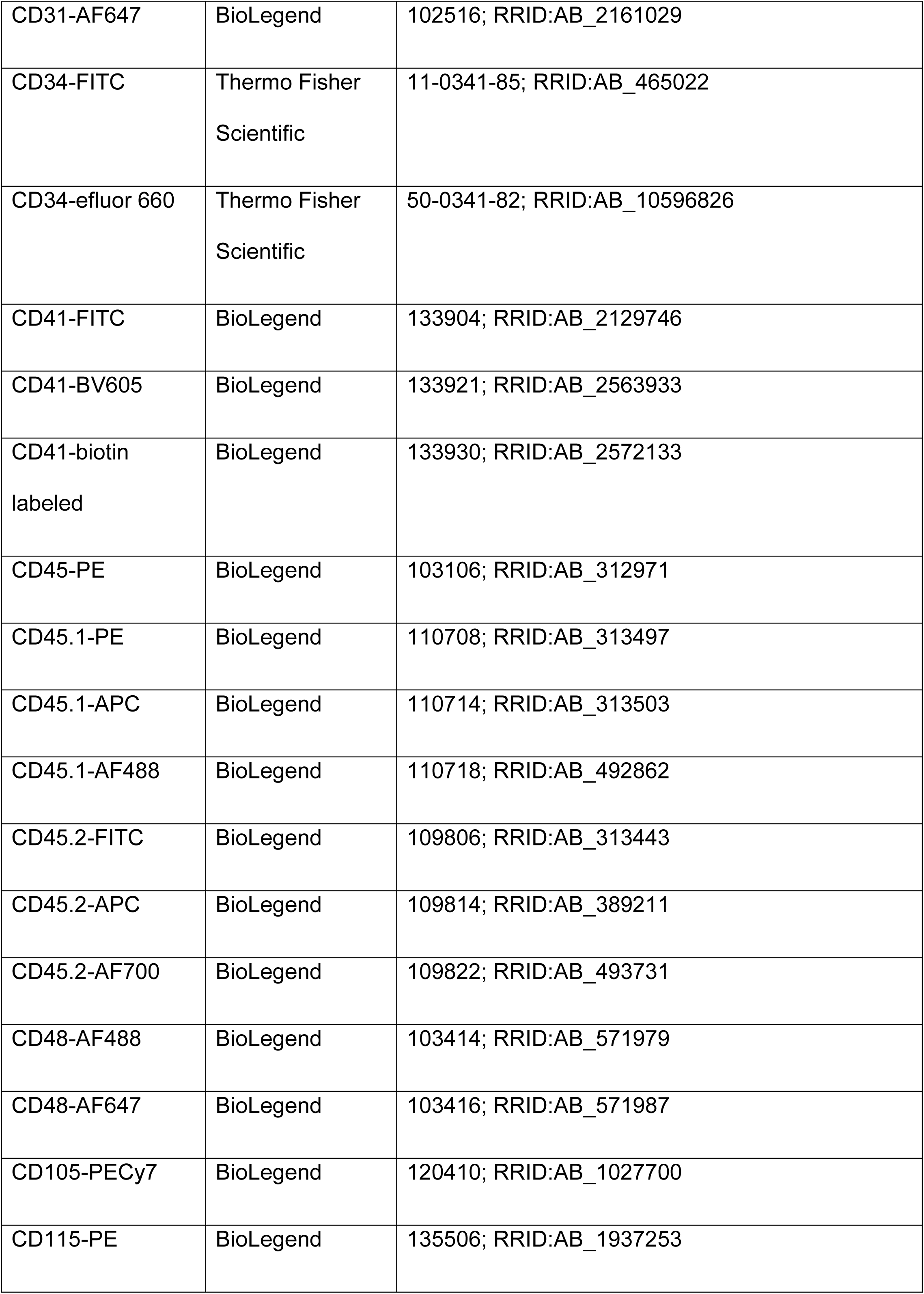
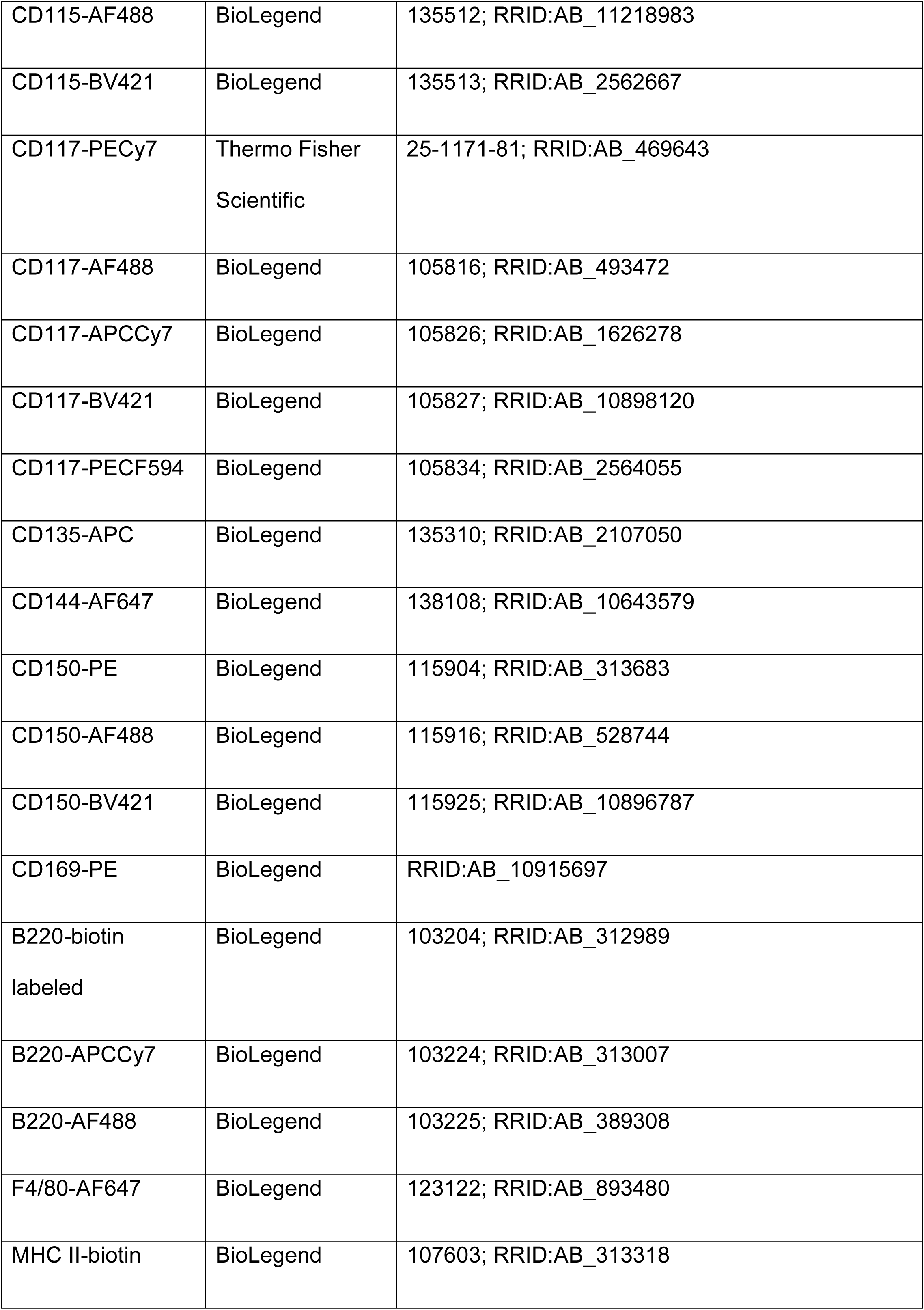
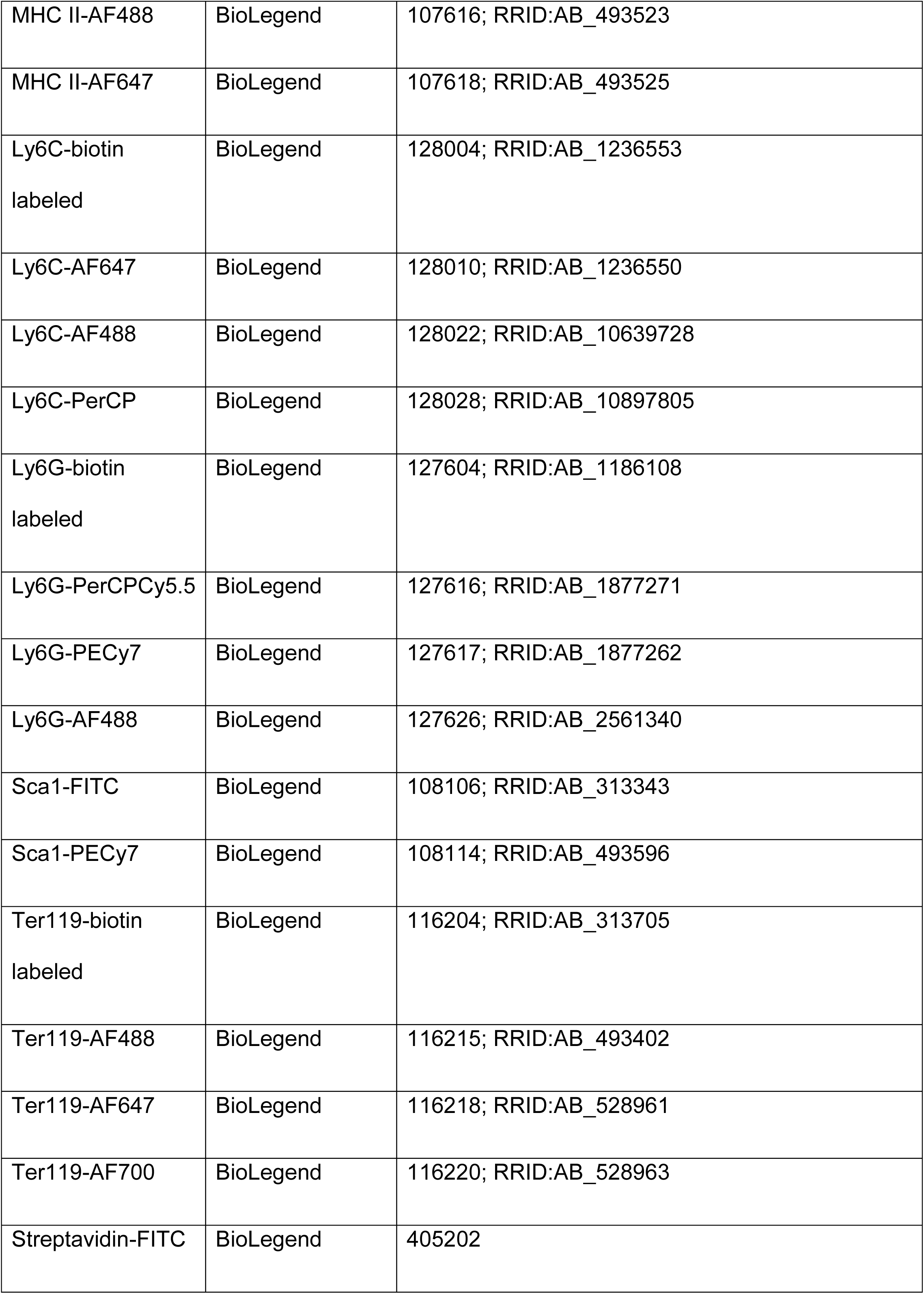
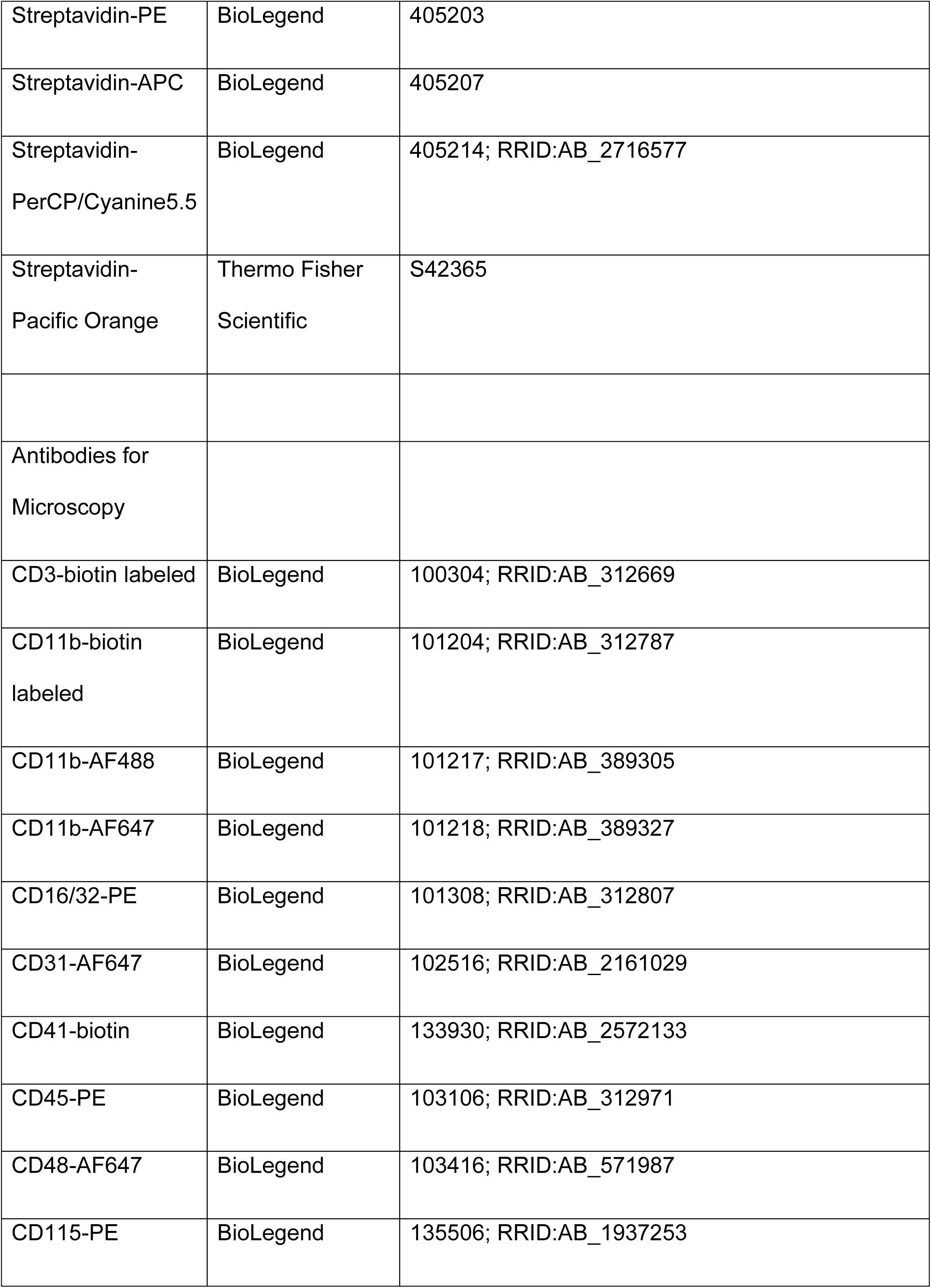
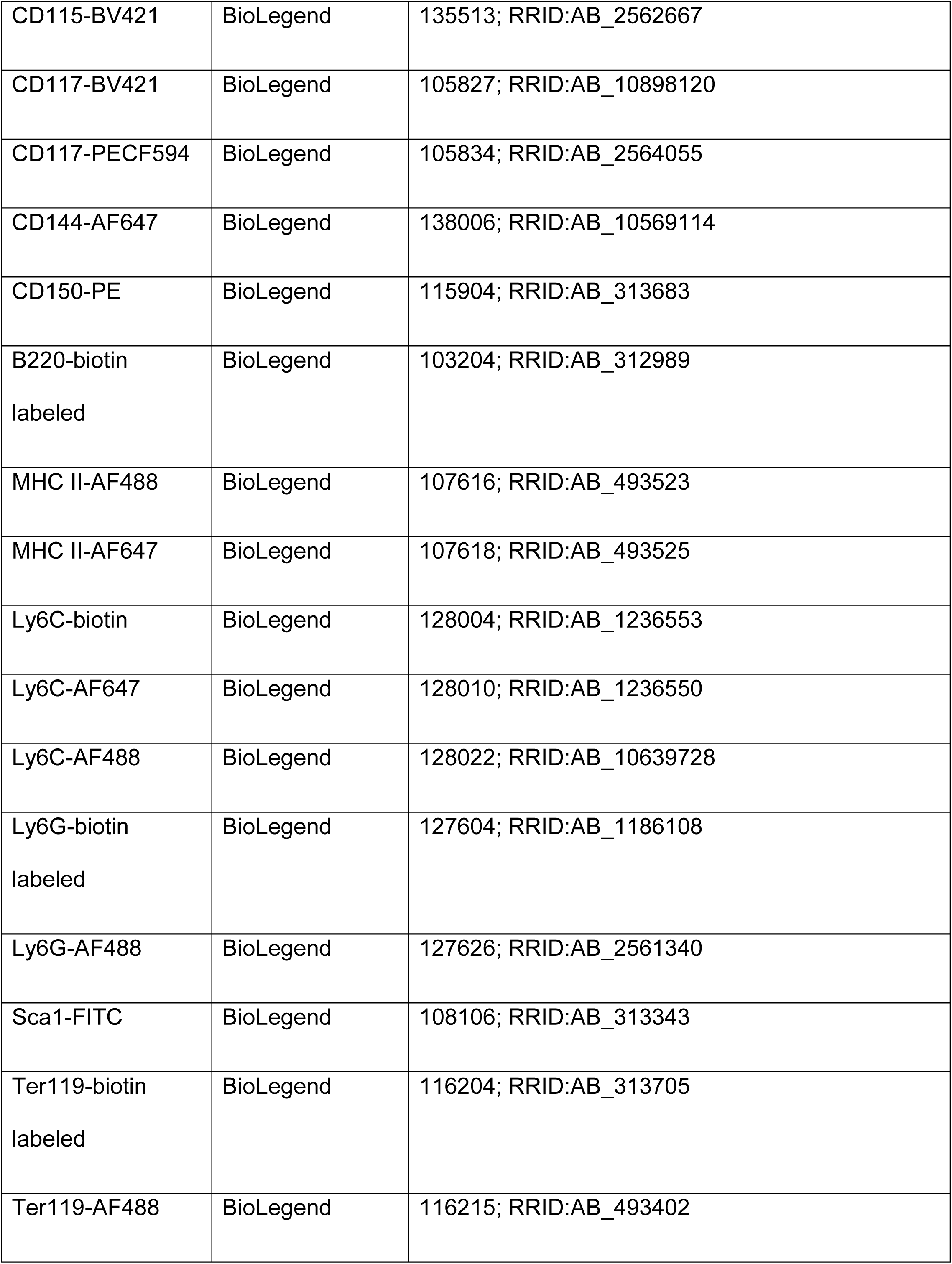
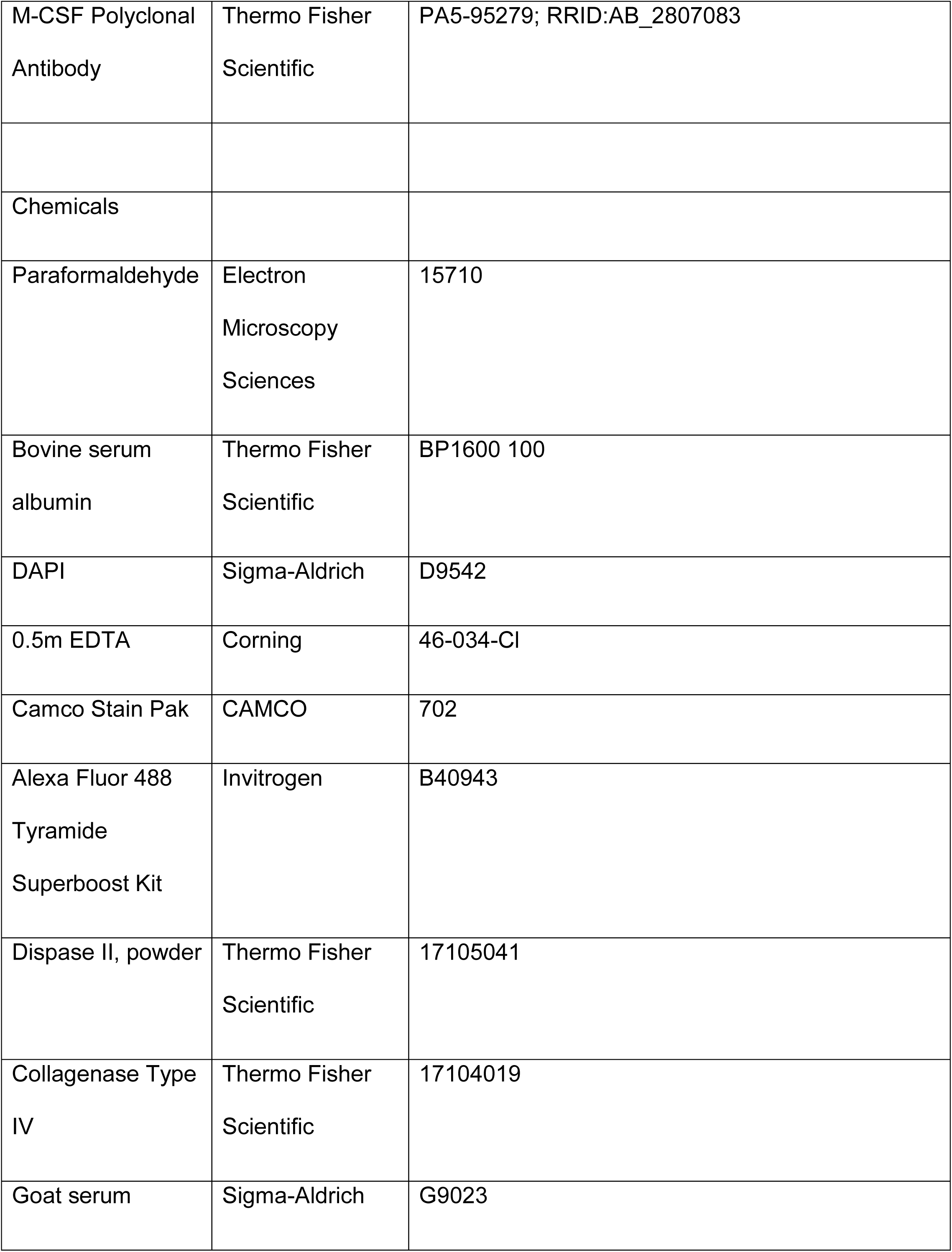
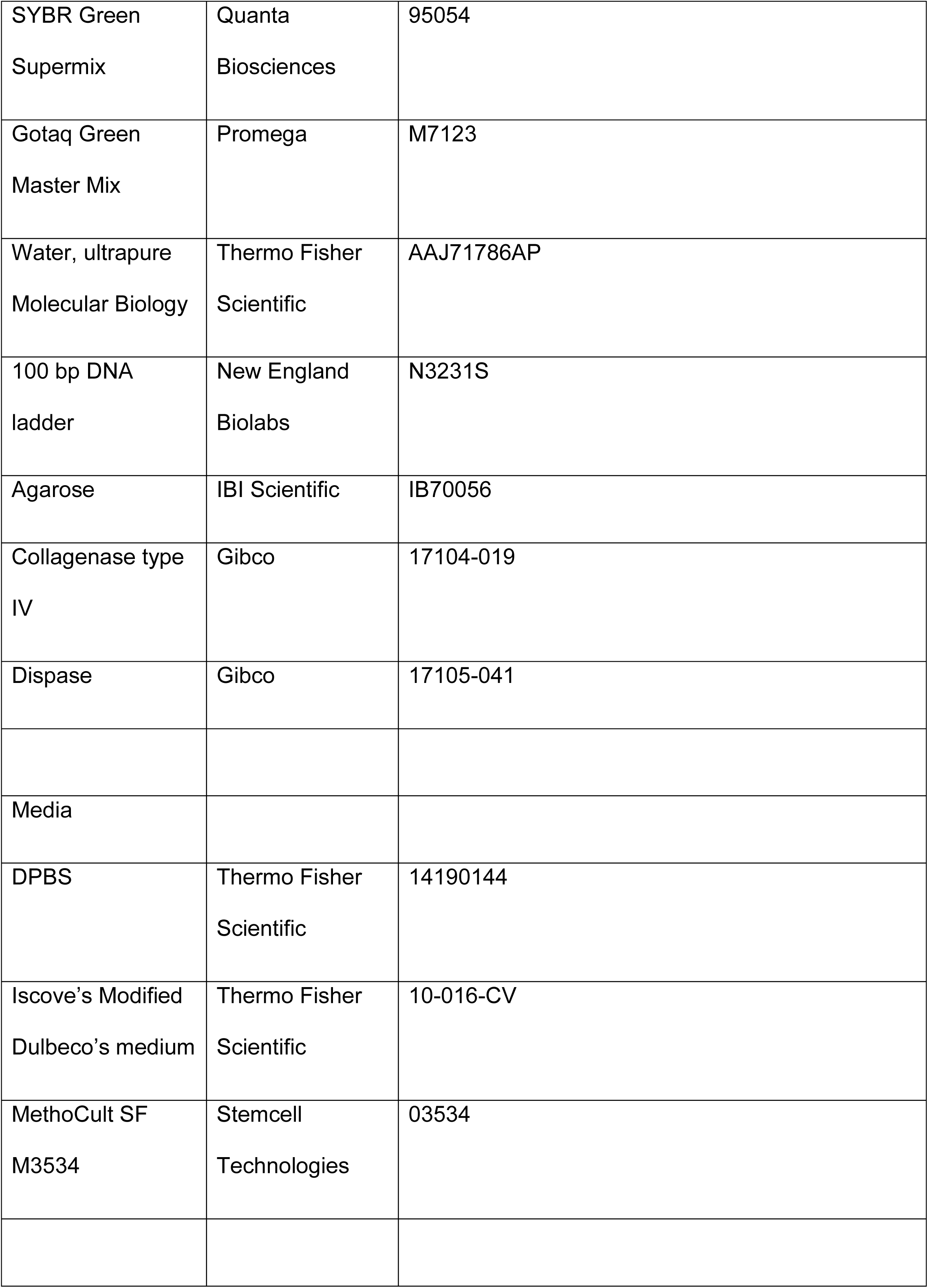
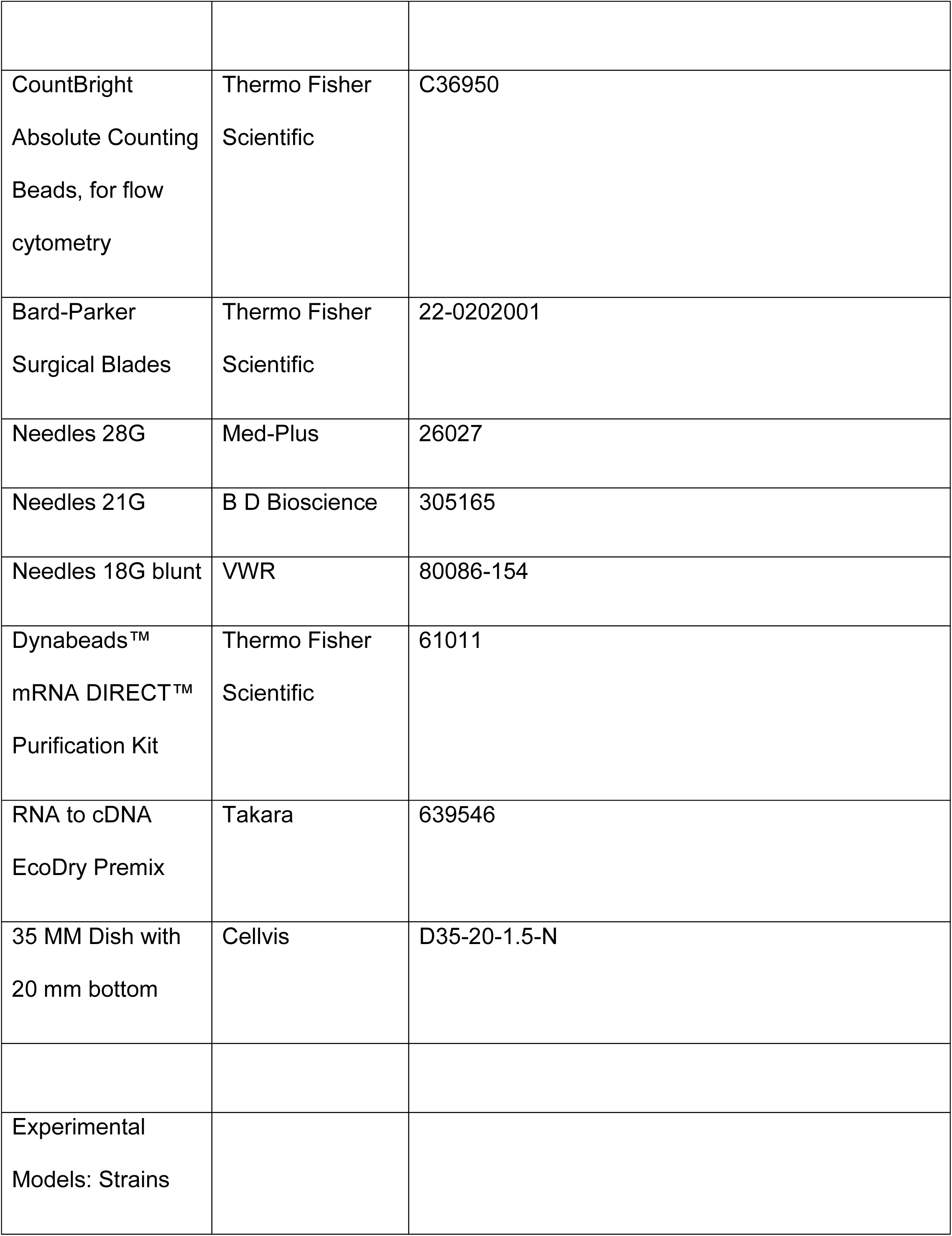
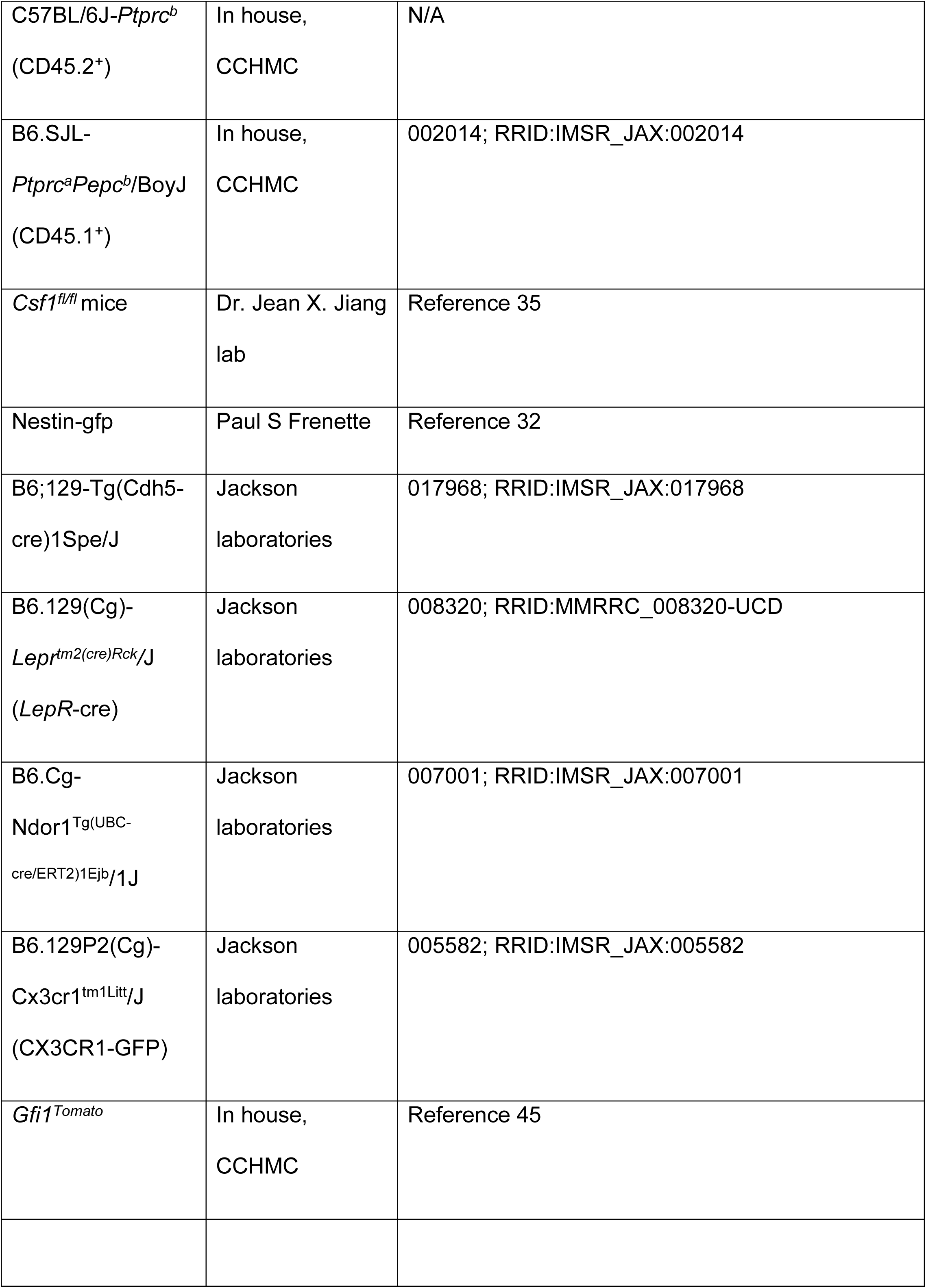
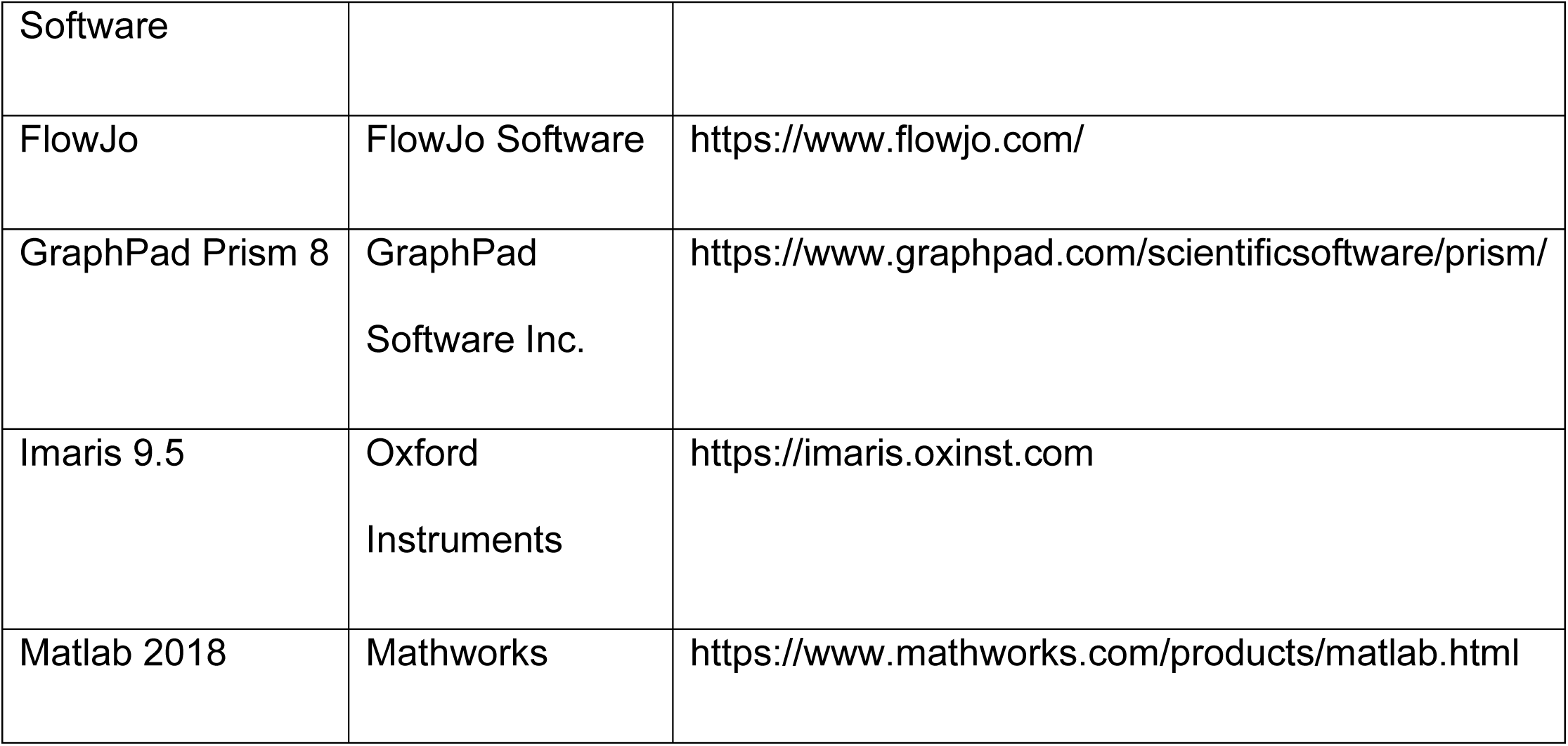

## Methods

### Mice

C57BL/6J*-Ptprc*^*b*^ (CD45.2^+^), B6.SJL-*Ptprc*^*a*^*Pepc*^*b*^/BoyJ (CD45.1^+^), B6;129-Tg(*Cdh5*-cre)1Spe/J, B6.129(Cg)-*Lepr*^*tm2(cre)Rck*^*/*J (*LepR*-cre), B6.Cg*-Ndor1*^*Tg(UBC-cre/ERT2)1Ejb*^*/1J* (UBC-cre/ERT2), and B6.129P2(Cg)*-Cx3cr1*^*tm1Litt*^/J (CX3CR1-GFP) mice were obtained from the Jackson Laboratory. *Nestin-gfp* mice^44^ were kindly provided by Paul. S. Frenette. *Csf1*^*fl/fl*^ and *Gfi1*^*Tomato*^ mice have been described previously^35,45^. *Csf1*^*+/-*^ mice were generated by breeding *Csf1*^*fl/fl*^ mice with UBC-cre/ERT2 mice^46^ and mating the offspring-after tamoxifen treatment-with C57BL/6J mice. We mated *Csf1*^*+/-*^ mice with *Cdh5*-cre^37^ and *LepR*-cre mice^36^ to generate *Cdh5*-cre:*Csf1*^*fl/-*^ mice and *LepR-*cre:*Csf1*^*fl/-*^ mice. Mice received water and food *ad libitum*. All experiments were performed in 8- to 14-week-old male and female mice. Mice were house at the vivarium at Cincinnati Children’s Hospital Medical Center under a 14h light: 10h darkness schedule. All studies were approved by the Animal Care Committee of Cincinnati Children’s Hospital Medical Center.

### Bone marrow and peripheral-blood collection for FACS analyses

Mice were euthanized by isoflurane inhalation followed by cervical dislocation. Bone marrow cells were harvested by flushing bones with 1 ml of ice-cold PEB buffer (2 mM EDTA and 0.5% bovine serum albumin in PBS). Blood was collected from the retro-orbital venous sinus in tubes containing EDTA. Red blood cells in peripheral blood were lysed by the addition of 1 ml of RBC lysis buffer (150 mM NH4Cl, 10 mM NaCO3 and 0.1 mM EDTA). Cells were immediately decanted by centrifugation, resuspended in ice-cold PEB and used in subsequent assays. We used CountBright™ Absolute Counting Beads (cat. no. C36950, Thermo Fisher Scientific) to count bone marrow and blood cell numbers in a flow cytometer according to the manufacturer’s protocol.

### Isolation of bone marrow stromal cells

Bone marrow digestion was performed as previously described^47^.

### FACS analyses

Cells were stained under dark for 30 min in PEB buffer containing antibodies, washed thrice with ice cold PBS and analyzed in a BD LSR II (BD Biosciences) instrument or FACS-purified with a BD FACS Aria II or a SH800S (Sony) cell sorter. Dead cells and doublets were excluded on the basis of FSC and SSC distribution and DAPI (Sigma) exclusion. Antibodies used were: B220 (clone RA3-6B2), CD3 (clone 145-2C11), CD8 (clone 53-6.7), CD11b (clone M1/70), CD11c (clone N418), CD16/32 (clone 93), CD31 (clone A20), CD41 (clone MWReg30), CD45 (clone 30-F11), CD45.1 (clone A20), CD45.2 (clone 104), CD48 (clone HM48-1), CD105 (clone MJ7/18), CD115 (clone AFS98), CD135 (clone A2F10), CD144 (clone BV13), CD150 (clone TC15-12F12.2), CD169 (clone 3D6.112), F4/80 (clone BM8), Ly6C (clone HK1.4), Ly6-G (clone 1A8), Sca-1 (clone D7), Ter119 (clone TER-119), MHC II (clone M5/114.15.2), from BioLegend; CD34 (clone RAM34) and CD117 (clone 2B8), from BioLegend or Thermo Fisher Scientific. Data was analyzed with FlowJo (Tree Star).

### Cytospin preparation and analyses

FACS-purified cells were decanted in 30 slides using a Cytospin 4 Cytocentrifuge (Thermo Fisher Scientific) following manufacturer’s instructions. Slides were stained with Camco Stain Pak (Cambridge Diagnostic Inc) according to the manufacturer’s instruction. This kit gives results similar to a Wright-Giemsa stain. Slides were analyzed on Zeiss AX10 inverted microscope (Carl Zeiss, Oberkochen, Germany).

### Colony-forming unit (CFU) assay

The indicated number of FACS-purified cells were seeded into methylcellulose culture medium (M3534, StemCell Technologies), plated in in 35 mm dishes and incubated at 37°C, in 5% CO2, with ≥ 95% humidity for 7-10 days. Colonies were identified and counted based on cluster size and cell morphology using Nikon Eclipse Ti inverted microscope (Nikon Instruments Inc., NY, USA).

### RNA isolation and qPCR analyses

RNA isolation was performed with a Dynabeads mRNA Direct Kit (Thermo Fisher Scientific, no. 61011) according to the manufacturer’s protocol. cDNA synthesis was performed with RNA to cDNA EcoDry Premix (Takara, 639546) according to the manufacturer’s instructions. qPCR was performed using SYBR Green Supermix (Quanta Biosciences, no. 95054) in an ABI PRISM 7900HT Sequence Detection System (Applied Biosystems). Results were analyzed in SDS 2.4 software (Applied Biosystems). Primers were as follows: CSF1 forward, 5’ -ATGAGCAGGAGTATTGCCAAGG- 3’; CSF1 reverse, 5’ -TCCATTCCCAATCATGTGGCTA- 3’; IRF8 forward, 5’ - AGACCATGTTCCGTATCCCCT- 3’, IRF8 reverse, 5’ -CACAGCGTAACCTCGTCTTCC −3’; GFI1 forward, 5’ -AGAAGGCGCACAGCTATCAC- 3’, GFI1 reverse, 5’ -GGCTCCATTTTCGACTCGC- 3’; mGAPDH forward, 5’ -TGTGTCCGTCGTGGATCTGA- 3’; mGAPDH reverse, 5’- CCTGCTTCACCACCTTCTTGA-3’.

### Competitive reconstitution assays in irradiated mice

Recipient mice were lethally conditioned with total 1175 rads irradiation dose (700 + 475 split dose 3 hours apart). 1 × 10^6^ bone marrow cells of control or conditional knockout mice were mixed with 1 × 10^6^ CD45.1^+^ competitor mouse bone marrow cells and transplanted by injection into the tail vein of CD45.1^+^ recipients.

### Whole-mount immunostaining

In some cases we injected retro-orbitally with 10 μg of Alexa Fluor 647 anti-mouse CD31 (BioLegend, no. 110724) and 10 μg of Alexa Fluor 647 anti-mouse CD144 (BioLegend, no. 138006) ten minutes before euthanasia to visualize the BM vasculature as described^32,48^.

Whole-mount sternum stain has been described before^49^. Sterna were processed immediately after euthanasia. After dissection we removed all connective tissue by gentle scraping with a blade. Fragments with bone marrow cavity were dissected and sectioned along the sagittal plane to expose the BM as described. Each half of the sterna was fixed in 4% PFA (Electron Microscopy Sciences, no. 15710) in DPBS for 3 hours. Each fragment was washed thrice with DPBS, followed by 1 hour blocking in DPBS containing 10% (v/v) goat serum (Sigma-Aldrich, no. G9023). We stained each sample with 100 µl staining buffer (2% goat serum in DPBS and the indicated antibodies). All steps were performed at 4°C.

### CSF1 detection

After fixation and blocking as above, sterna were stained with 100 µl of staining buffer containing anti CSF1 (Thermo Fisher Scientific, no. PA5-95279, 1:200 dilution), anti MHCII (BioLegend, no. 107618) and 2% goat serum in DPBS at 4°C for 12h. Each sternum was washed thrice with DPBS and then incubated with Superboost HRP conjugated anti-rabbit antibody (Invitrogen, no. B40943) in DPBS at room temperature for 1 hour. After washing three times each sternum was developed by incubation in 100 µl of tyramide solution (Invitrogen, no. B40943) for 8 minutes. Sterna were washed three additional times in DPBS and used for confocal analyses.

### Confocal imaging

We used a Nikon A1R GaAsP Multiphoton Upright Confocal Microscope and a Nikon A1R GaAsP Inverted Microscope. Specifications for the Upright Confocal: 405 nm, 488 nm, 561 nm, and 638 nm diode lasers, Coherent Chameleon II TiSapphire IR laser, tunable from 700-1000 nm, fully encoded scanning XY motorized stage, Piezo-Z nosepiece for high-speed Z-stack acquisition (100 μm/s), resonant and galvanometric scanners, four high-quantum efficiency, low-noise Hamamatsu photomultiplier tubes, a transmitted PMT for transmitted light for 400-820 nm detection, four high-quantum efficiency GaAsP non-descanned detectors for multiphoton imaging. We used a 16X Apo 0.8 NA LWD Water Immersion Objective with a 3.0 mm working distance. Images were taken using the resonant scanner with 8X line averaging, 1024 × 1024 pixel resolution, 2 μm Z-step, and pinhole at 21.7 μm. Bone signal was got from second harmonic generation (SHG) imaging with 840 nm excitation.

Specifications for the Inverted Confocal: 405 nm, 442 nm, 488 nm, 561 nm, 640 nm, and 730 nm diode lasers, fully encoded scanning XY motorized stage, Piezo-Z stage insert for high-speed Z-stack acquisition (100 μm/s), resonant and galvanometric scanners, two high-quantum efficiency, low-noise Hamamatsu photomultiplier tubes, a transmitted PMT for transmitted light, and two very high-QE Gallium-Arsenide-Phosphide PMTs for overall 400-820 nm detection. We used a 40X Apo 1.15 NA LWD DIC-Water Immersion Objective with a 0.59 mm working distance for high power image. Images were taken using the resonant scanner with 8X line averaging, 1024 × 1024 pixel resolution, 0.5 μm Z-step, and pinhole at 50 μm.

### Image and distance analyses

We used Nikon Elements software (5.20.02), Imaris x64 software (9.3) and Matlab software (2018a) installed in a HP Z4 64-bit workstation with Dual Intel Xeon Processor W-2145, 64GB ECC-RAM, 8 GB NVIDA Quadro RTX5000 Graphics card for all our analyses. We only analyzed images for which the whole sternum fragment (6-8 images) was successfully imaged. These images were stitched together using Nikon Elements. Stitched whole bone images were further processed by an artificial intelligence algorithm (Denoise.AI) for noise removal using Nikon elements. We used Imaris to identify each cell, replace it with a color-coded sphere and obtain its X, Y and Z coordinates. We also used Imaris to create surfaces for sinusoids, arterioles and the endosteal surface. For each cell type we measured the diameter of 50-150 cells of each type to obtain the mean diameter (HSC=8.84±1.56 μm; GMP=12.68±1.52 μm, MDP=12.13±1.19 μm, GP=11.70±0.99 μm, MOP=11.49±1.25 μm, PN=10.21±1.08 μm, IN=8.72±0.75 μm, MN=8.10±1.09 μm, Ly6C^hi^ Mo=9.25±0.86 μm, Ly6C^lo^ Mo=9.30±1.17 μm, cDC=12.33±2.69 μm). Since cDC have reticular shapes we measured the cell bodies to obtain the median diameter. We used Imaris to measure the distance from each cell to the closest vascular structure or the endosteum and then subtracted the mean radius for each cell type. To quantify cell to cell distance we exported the coordinates of the cells of interest to Matlab and then used an algorithm to quantify the distance from the center of each cell to the center of all other cells. We then subtracted the mean radius of each cell from these numbers. The graph summarizing the distances between the cell populations examined in this manuscript was assembled using Cytoscape using Prefuse force directed layout with edge distances weighted based on the median experimentally quantified cell-type interactions length^50^.

Note that in all the images in the manuscript we adjusted the brightness and contrast of each channel to be able to detect negative, dim and bright cells for each fluorescent signal.

### Random simulations

We stained and imaged a sternum fragments with CD45 and Ter119 antibodies to detect hematopoietic cells; CD31, CD144 and Sca1 antibodies to detect sinusoids and arterioles; and used second harmonic generation to detect bone. These images were processed as above to obtain the coordinates of all hematopoietic cells (59,659 cells), vessels and bone in the sternum. We then used Research Randomizer^51^ to randomly select dots representing each type of myeloid cell at the same frequencies found in vivo through the bone marrow cavity and measured the distances between these random cells or with vessels and bone as above. Each random simulation was repeated 100-200 times.

### Stromal UMAP analysis

To identify diverse stromal, hematopoietic and other cell populations we reanalyzed 19 independent 10x Genomics captures (GSE128423^34^). The Cell Ranger produced filtered sparse matrics outputs. The merged counts files from these data were scaled and normalized in the software AltAnalyze (CountsNormalize function). We identified 46 preliminary transcriptionally distinct cell populations in 89,007 cells based on unsupervised using the software Iterative Clustering and Guide-gene Selection (ICGS) version 2. To annotate these populations and identify sub-clusters based on prior knowledge, we performed a secondary analysis using the supervised classification tool cellHarmony, comparing all cells to reference hematopoietic (GSE120409^52^) and sorted stromal populations (GSE108891^53^), resulting in 61 final annotated cell populations (Supplemental Table 2). Visualization of clusters and marker genes was performed using UMAP visualization in AltAnalyze.

### Statistics

For graphs quantifying cells in different mice we indicate the mean and each dot corresponds to one mouse. For graphs showing distances between cells and structures each dot corresponds to one cell and the horizontal bar indicates the median. Statistical differences were calculated using two-tailed Student’s T tests. *P < 0.05; **P < 0.01; ***P < 0.001; ****P < 0.0001; NS, not significant.

### Data reporting

No statistical methods were used to predetermine sample size. All mice were included in the analyses. Mice were randomly allocated to the different groups on the basis of cage, genotype, and litter size. For all experiments, we aimed to have the same number of mice in the control and experimental groups. Investigators were not blinded to allocation during experiments and outcome assessment.

## Acknowledgements

We would like to thank Drs. Jose Cancelas, Marie-Dominique Filippi, Damien Reynaud, Daniel Starczynowski, and Andres Hidalgo for valuable feedback on the manuscript; the Confocal Imaging Core, the Research Flow Cytometry Core and the Veterinary Services at the University of Michigan and Cincinnati Children’s Medical Center for excellent experimental and technical assistance. This work was partially supported by the National Heart Lung and Blood Institute (R01HL122661 to H.L.G, and R01HL136529 to D.L.).A. S. is supported by T32 AI118697/AI/NIAID NIH HHS/United States Data was generated using the SH800 cell sorter funded by NIH S10OD023410.

## Author contributions

D.L conceptualized and managed the study. D.L., J.Z., H.L.G., N.S., J.D.E., J.M.K, A.D., and Q.W. designed experiments. J.Z. and Q.W. developed all the stains to analyze myelopoiesis in situ. J.Z., Q.W., C.B.J., A.O., A.S., M.M., B.W., and E. B., performed and analyzed experiments. J.X.J and S.L.A. generated the *Csf1*^*fl/fl*^ mice. N.S. and L.H. performed bioinformatics analyses. D.L. and J.Z., assembled the figures and wrote the manuscript with input from all the authors.

## Additional information

**Supplementary information** is available for this paper. Correspondence and requests for materials should be addressed to Daniel Lucas.

## Data availability

Source Data for quantifications described in the text or shown in graphs plotted in Figs. 1–4 and Extended Data Figs. 1–5 are available with the manuscript.

**Three dimensional mapping identifies distinct vascular niches for myelopoiesis**

