## Extended Data Figures 1-5 and Supplemental Table 1 for "Three-dimensional mapping identifies distinct vascular niches for myelopoiesis"

**Extended data legends**

**Supplementary Table 1.** Table indicating how each cell type was identified in the different stains shown in Fig. 1a.

**Supplementary Table 2.** Table showing the different ICGS2 marker genes and Cell barcode assignments for the difference cell clusters identified from the reanalyses of 19 independent 10x Genomics captures (GSE128423[34]).

**Extended Data Figure 1. a.** FACS plots showing the gating strategy to identify MDP and MOP (as described in reference[26]). **b.** Gating strategy to identify GMP, GP, and MOP (as described in reference[24]). The Lineage panel contains antibodies against Ly6G, CD11b, Ter119, B220 and CD3. **c.** Images showing that Ly6C labels CD31$^+$CD144$^+$Sca1$^+$ arterioles; and quantification showing that all Sca1$^+$ arterioles are also Ly6C$^+$. Scale bars are 200µm. **d, e.** FACS plots (d) showing that only the CD11b$^+$ gate contains CD117$^+$CD115$^+$ or CD117$^+$Ly6C$^+$ cells and histograms (e) showing that GP and MOP are the only Ly6C$^+$ cells in the Lin$^-$CD117$^+$ gate. Together these data indicate that CD11b alone can be used to replace the Lineage panel to exclude contamination of mature cells when detecting MDP, MOP and GP. **f**. Representative image showing that CD11b$^-$CD117$^+$CD115$^+$Ly6C$^-$ MDP and CD11b$^-$CD117$^+$CD115$^+$Ly6C$^+$ MOP are GFP$^+$ in *Cx3cr1-gfp* mice. Scale bar = 10µm. **g**. qPCR showing *Gfi* and *Irf8* expression (relative to *Gapdh*) in FACS-purified GP or MOP. **h**. FACS analyses in *Gfi-tdTomato* mice showing differential tdTomato expression in GP and MOP. **i**. Quantification of promyelocytes (PM), myelocytes (MC), metamylocytes (MM), banded cells (BC) and polymorphonucleated neutrophils (PMN) in cytospin preparations of FACS purified Pre-Neutrophils, Immature Neutrophils and Mature Neutrophils. N = 2 mice. Scale bar = 10µm. **j**. The image shows Cx3cr1-GFP$^+$MHCII$^+$ dendritic cells in the bone marrow and the panels show that all MHCII$^+$ reticular cells are $CD11b^+$ and Cx3cr1-$GFP^+$, n = 3 mice. **k**. Image showing that Cx3cr1-$GFP^+MHCII^+$ dendritic cells are also $CD11b^+$ but do not express B220 or CD8. Scale bar = 10μm. In all bar graphs one dot corresponds to one mouse.

**Extended Data Figure 2. a.** Scheme showing the experimental pipeline to identify and color code myeloid cells in the bone marrow and to generate random distributions. Scale bar = 200μm. **b.** Histograms showing the observed distribution of distances from each GMP (blue), MDP (green), MOP (orange) and GP (red) or random cells (white) to the closest indicated cell (n = 86 GMP from 4 sterna of 4 mice, n = 243 MDP from 23 sterna of 15 mice, n = 458 MOP from 11 sterna of 11 mice, n = 338 GP from 15 sterna of 12 mice). **c.** Representative image showing a Pre-neutrophil cluster lacking GP and the histograms showing the observed distribution of distances from each Pre-Neutrophil to the six closest Pre-Neutrophils (upper panel) or Immature Neutrophils (lower panels) in clusters with or without GP (n = 2493 Pre-Neutrophil from 3 sterna of 3 mice). Scale bar = 10μm. **d-i.** Histograms showing the observed (color) and random (white) distribution of distances from each $Ly6C^{lo}$ Mo (yellow), MDP (green), MOP (orange) and cDC (pink) to the six closest indicated cells d**.** n = 1603 $Ly6C^{lo}$ monocytes from 3 sterna of 3 mice; e**.** n = 67 MDP from 6 sterna of 4 mice; f**.** n = 171 MOP from 4 sterna of 4 mice; g**.** n = 1228 cDC from 6 sterna of 6 mice; h**.** n = 139 MDP from 11 sterna of 6 mice; **i.** n = 200 MOP from 5 sterna of 4 mice).

**Extended Data Figure 3. a.** Representative image showing simultaneous detection of Pre- and Immature Neutrophils, $Ly6C^{lo}$ monocytes and cDC. Scale bar = 10μm. **b.** Representative image showing detection of HSC and MDP in a single stain. Scale bar = 10μm. **c.** Representative image showing detection of HSC and a population containing $CD117^+Ly6C^+$ GP and MOP in a single stain. Scale bar = 10μm. **d.** Map showing the location of the HSC and GP/MOP cells in the bone marrow. **e.** Histograms showing the distance from each $Lin^-CD117^+Ly6C^+$ cells (either GP or MOP, pink dots) or the random simulation (white dots) to the closest HSC. Each dot corresponds to one cell and the dot radius is three times the average cell radius (n = 191 GP and MOP from 3 sterna of 3 mice).

**Extended Data Figure 4. a.** qPCR analyses showing *Csf1* mRNA levels (relative to GAPDH) in FACS purified BM endothelial cells or Nestin-GFP$^{dim}$ perivascular cells (which largely overlap with LepR$^{+}$ perivascular cells[43]). **b.** qPCR analyses showing *Csf1* mRNA levels (normalized to Control) in FACS-purified CD45$^{-}$Ter119$^{-}$CD31$^{-}$LepR$^{+}$ cells or endothelial cells in control, $Csf1^{\Delta LepR}$, or $Csf1^{\Delta EC}$ mice. **c, d.** Number of BM cellularity or the indicated populations in the femur of control or $Csf1^{\Delta LepR}$ mice. **e.** Colony forming activity (d, black: CFU-GM, grey: CFU-G, white: CFU-M) of the indicated progenitors FACS-purified from control (n = 5) or $Csf1^{\Delta LepR}$ (n = 5) mice. **f.** Number of the indicated populations in the femur of control or $Csf1^{\Delta LepR}$ mice. **g.** Colony forming activity (black: CFU-GM, grey: CFU-G) of the indicated progenitors FACS-purified from control (n = 5) or $Csf1^{\Delta E}$ (n = 5) mice. **h, i.** Number (h) and CFU-M activity (i) of GMP or MOP from control or $Csf1^{\Delta EC}$ mice. **j-l.** Number of BM cellularity or the indicated populations in the femur or blood of control or $Csf1^{\Delta EC}$ mice**. m**. The panels show the percentage of the indicated CD45.2$^{+}$ cells in the blood of lethally irradiated CD45.1$^{+}$ recipients after transplant of $10^{6}$ BM cells purified from *Ctrl* ( white dots) or $Csf1^{\Delta EC}$ mice (red)- both CD45.2$^{+}$- together with $10^{6}$ CD45.1$^{+}$ competitor cells at the indicated time points after transplantation. The dots show the mean of 12 mice per group and the error bars show standard deviation. Unless otherwise indicated for all panels one dot corresponds to one mouse.

**Extended Data Figure 5. a.** Number of cDC found within the indicated distances of CSF1$^{+}$ and CSF1$^{-}$ vessels in wild-type (n = 76 CSF1$^{+}$ vessels and n = 520 CSF1$^{-}$ vessels in 4 sterna of 3 wild-type mice). **b.** Histograms showing the distance from each cDC to the closest sinusoid in control or $Csf1^{\Delta EC}$ mice (n= 451 cDC in 2 sterna of control mice, n = 343 cDC in 3 sterna of $Csf1^{\Delta EC}$ mice). **c.** Maps showing the relocation of MDP away from sinusoids in $Csf1^{\Delta EC}$ mice. Scale bars = 200 and 10µm. The radius of the dots is 3x (left map) or 1x (right images) the average radius of the MDP. **d.** Histograms showing the distribution of distances from each MDP to the six closest Ly6C$^{lo}$ Mo or cDC in control or $Csf1^{\Delta EC}$ mice (For MDP-Ly6C$^{lo}$ monocyte, n = 37 MDP from 4 sterna of 3 Control mice, n

= 18 MDP from 4 sterna of 3 *Csf1*$^{ΔEC}$. For MDP-cDC, n = 47 MDP from 6 sterna of 3 Control mice, n = 47 MDP from 9 sterna of 3 *Csf1*$^{ΔEC}$).

**Supplementary Video 1.** Representative video showing how the we identified and annotated the different cell populations. Bone marrow cells were stained with CD11b (blue),
CD115 (red), CD117(magenta), Ly6C (green), and Ly6G (white). MOP were identified as CD11b$^{-}$CD117$^{+}$CD115$^{+}$Ly6C$^{+}$ cells; Ly6C$^{hi}$ Mo were identified as CD11b$^{+}$CD117$^{-}$CD115$^{+}$Ly6C$^{hi}$ cells; Ly6C$^{lo}$ Mo were identified as CD11b$^{+}$CD117$^{-}$CD115$^{+}$Ly6C$^{lo}$ cells; MDP were identified as CD11b$^{-}$CD117$^{+}$CD115$^{+}$Ly6C$^{-}$ cells; GP were identified as CD11b$^{-}$CD117$^{+}$CD115$^{-}$Ly6C$^{+}$ cells; Pre neutrophil (PN) were identified as CD11b$^{+}$CD117$^{dim}$CD115$^{-}$Ly6G$^{lo}$ cells; Immature neutrophil (IN) were identified as CD11b$^{+}$CD117$^{dim}$CD115$^{-}$Ly6G$^{hi}$ cells; and Mature neutrophil (MN) were identified as CD11b$^{+}$CD117$^{-}$CD115$^{-}$Ly6G$^{hi}$ cells, We used Imaris software to identify each cell, replace it with a color-coded sphere (dot), and obtain its X, Y and Z coordinates. Each dot radius corresponds to the average radius of the replaced cell.

**Supplementary Video 2.** Representative video showing the spatial distribution of GMP, MDP, MOP, and GP in a mouse sternum. The position of each progenitor is indicated by a color-coded dot. Note that the radius of each dot is 3x the average radius of the replaced cell.

**Supplementary Video 3.** Representative video showing the spatial distribution of GP, pre-neutrophils (PN), immature neutrophils (IN), and mature neutrophils (MN) in a mouse sternum. The position of each type of cell is indicated by a color-coded dot. The radius of each dot is 3x the radius of GP, 2x the radius of PN and In, and 1.5x the radius of MN.

**Supplementary Video 4.** Representative video showing the spatial distribution of MDP, MOP, Ly6C$^{hi}$ monocyte, Ly6C$^{lo}$ monocyte, and cDC in a mouse sternum. The position of each type of cell is indicated by a color-coded dot. The radius of each dot is 3x the radius of MDP and MOP and 2x the radius of all other cells.

**Supplementary Video 5.** Representative video showing neutrophil differentiation around a GP. The position of each type of cell is indicated by a color-coded dot. The radius of each dot matches the radius of each cell.

**Supplementary Video 6.** Representative video showing Ly6C$^{lo}$ Mo and cDC localization near MDP. The position of each type of cell is indicated by a color-coded dot. The radius of each dot matches the radius of each cell.

**Supplementary Video 7.** Representative video showing the interaction between MDP and sinusoids. The position of the MDP is denoted by a green dot, and the surface of the sinusoid was digitally reconstructed based on the signals for CD31 and CD144.

**Supplementary Video 8.** Representative video showing the interaction between MOP and sinusoids. The position of the MOP is denoted by an orange dot, and the surface of the sinusoid was digitally reconstructed based on the signals for CD31 and CD144.

**Supplementary Video 9.** Representative video showing the interaction between GP and sinusoids. The position of the GP is denoted by a red dot, and the surface of the sinusoid was digitally reconstructed based on the signals for CD31 and CD144.

**Supplementary Video 10.** Representative video showing the interaction between cDC and CSF1$^{+}$ sinusoids. Bone marrow cells were stained with antibodies against CD31 and CD144 (white), CSF1 (green), and MHC II (magenta). The position of each cDC is denoted by a magenta dot; the surface of the sinusoid was digitally reconstructed based on the signals for CD31 and CD144.  $CSF1^-$ vessels are shown in blue and $CSF1^+$ vessels in yellow.

| | **Stain 1** | **Stain 2** | **Stain 3** |
|---|---|---|---|
| **GMP** | Lin$^-$CD117$^+$CD115$^-$Ly6C$^-$CD16/32$^+$ | NA | NA |
| **MDP** | Lin$^-$CD117$^+$CD115$^+$Ly6C$^-$ | CD11b$^-$CD117$^+$CD115$^+$Ly6C$^-$ | CD11b$^-$CD117$^+$CD115$^+$Ly6C$^-$ |
| **MOP** | Lin$^-$CD117$^+$CD115$^+$Ly6C$^+$CD16/32$^+$ | CD11b$^-$CD117$^+$CD115$^+$Ly6C$^+$ | CD11b$^-$CD117$^+$CD115$^+$Ly6C$^+$ |
| **GP** | Lin$^-$CD117$^+$CD115$^-$Ly6C$^+$CD16/32$^+$ | CD11b$^-$CD117$^+$CD115$^-$Ly6C$^+$ | CD11b$^-$CD117$^+$CD115$^-$Ly6C$^+$ |
| **Ly6Chi Mo** | CD117$^-$CD115$^+$Ly6C$^{hi}$ | CD11b$^+$CD117$^-$CD115$^+$Ly6C$^{hi}$ | CD11b$^+$CD117$^-$CD115$^+$Ly6C$^{hi}$ |
| **Ly6Clo Mo** | CD117$^-$CD115$^+$Ly6C$^{lo}$ | CD11b$^+$CD117$^-$CD115$^+$Ly6C$^{lo}$ | CD11b$^+$CD117$^-$CD115$^+$Ly6C$^{lo}$ |
| **PN** | NA | CD11b$^+$CD117$^{dim}$CD115$^-$Ly6G$^{lo}$ | CD11b$^+$CD117$^{dim}$CD115$^-$ |
| **IN** | NA | CD11b$^+$CD117$^{dim}$CD115$^-$Ly6G$^{hi}$ | |
| **MN** | NA | CD11b$^+$CD117$^-$CD115$^-$Ly6G$^{hi}$ | NA |
| **cDC** | NA | NA | MHCII$^+$ reticular cells |

**Extended Data Table 1.** Table indicating how each cell type was identified in the different stains shown in Fig. 1a.

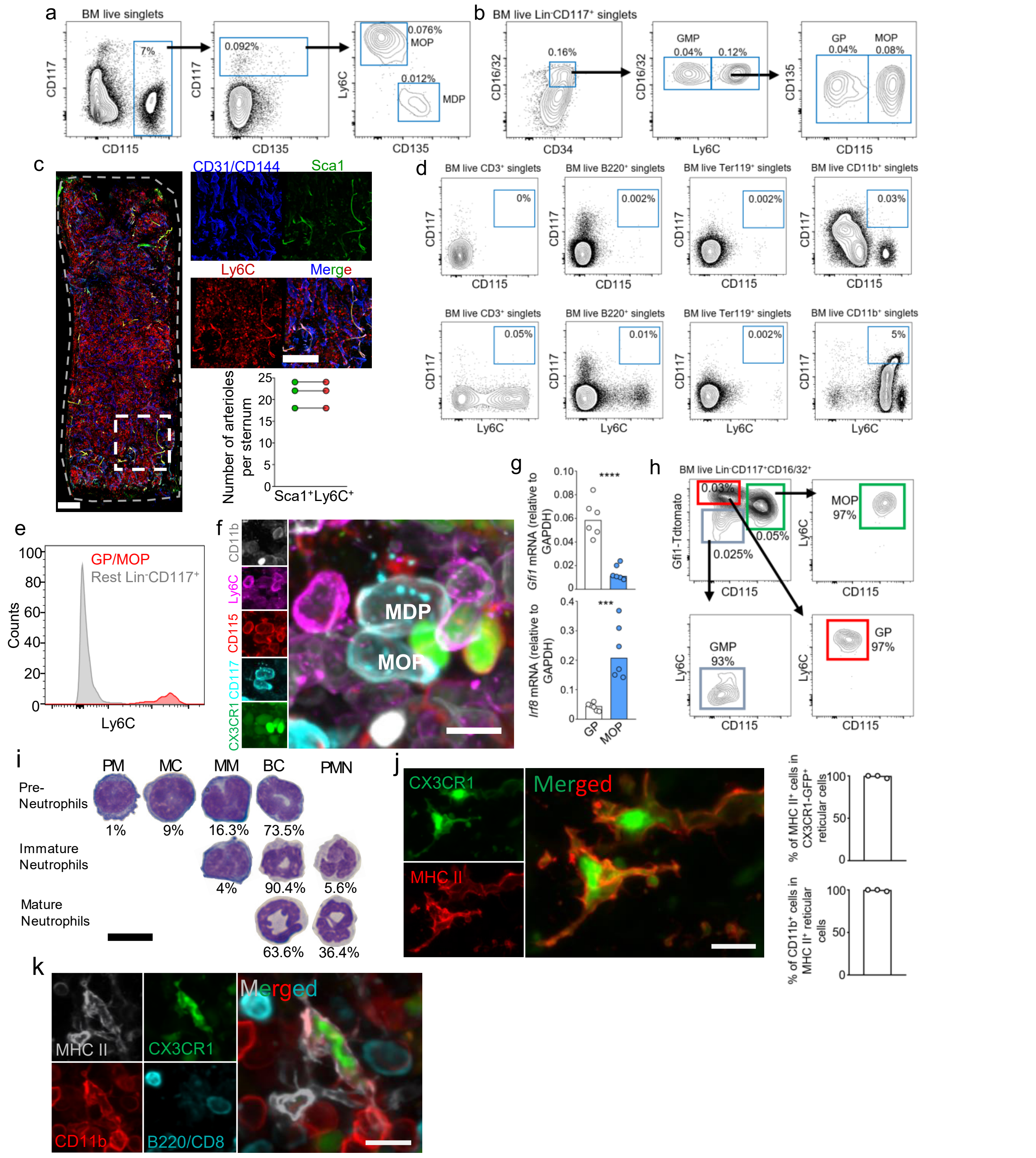


**Extended Data Figure 1. a.** FACS plots showing the gating strategy to identify MDP and MOP (as described in reference[26]). **b.** Gating strategy to identify GMP, GP, and MOP (as described in reference[24]). The Lineage panel contains antibodies against Ly6G, CD11b, Ter119, B220 and CD3. **c.** Images showing that Ly6C labels CD31$^+$CD144$^+$Sca1$^+$ arterioles; and quantification showing that all Sca1$^+$ arterioles are also Ly6C$^+$. Scale bars are 200μm. **d, e.** FACS plots (d) showing that only the CD11b$^+$ gate contains CD117$^+$CD115$^+$ or CD117$^+$Ly6C$^+$ cells and histograms (e) showing that GP and MOP are the only Ly6C$^+$ cells in the Lin$^-$CD117$^+$ gate. Together these data indicate that CD11b alone can be used to replace the Lineage panel to exclude contamination of mature cells when detecting MDP, MOP and GP. **f.** Representative image showing that CD11b$^-$CD117$^+$CD115$^+$Ly6C$^-$ MDP and CD11b$^-$CD117$^+$CD115$^+$Ly6C$^+$ MOP are GFP$^+$ in *Cx3cr1-gfp* mice. Scale bar = 10μm. **g.** qPCR showing *Gfi* and *Irf8* expression (relative to *Gapdh*) in FACS-purified GP or MOP. **h.** FACS analyses in *Gfi-tdTomato* mice showing differential tdTomato expression in GP and MOP. **i.** Quantification of promyelocytes (PM), myelocytes (MC), metamylocytes (MM), banded cells (BC) and polymorphonucleated neutrophils (PMN) in cytospin preparations of FACS purified Pre-Neutrophils, Immature Neutrophils and Mature Neutrophils. N = 2 mice. Scale bar = 10μm. **j.** The image shows Cx3cr1-GFP$^+$MHCII$^+$ dendritic cells in the bone marrow and the panels show that all MHCII$^+$ reticular cells are CD11b$^+$ and Cx3cr1-GFP$^+$, n = 3 mice. **k.** Image showing that Cx3cr1-GFP$^+$MHCII$^+$ dendritic cells are also CD11b$^+$ but do not express B220 or CD8. Scale bar = 10μm. In all bar graphs one dot corresponds to one mouse.

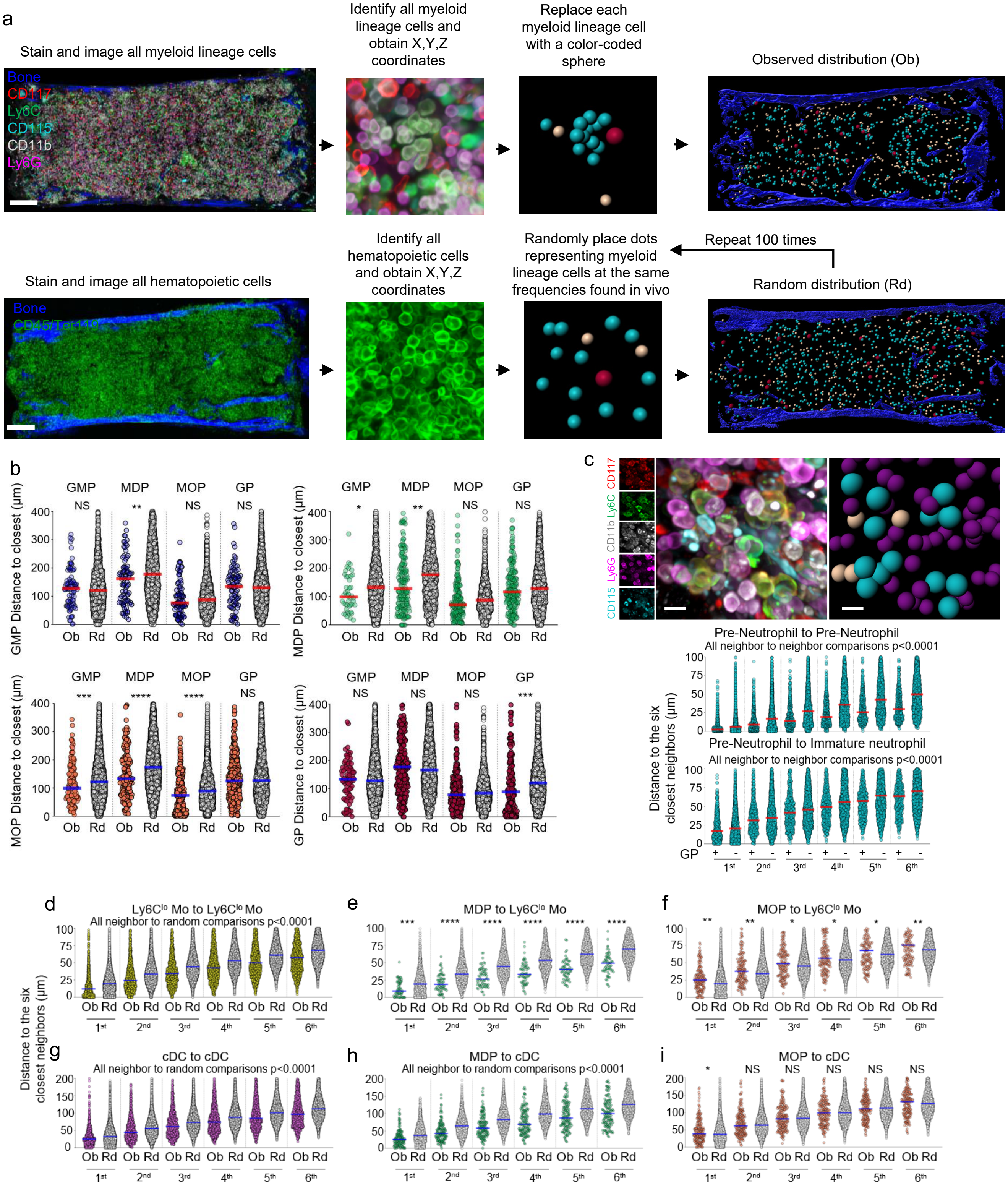


**Extended Data Figure 2. a.** Scheme showing the experimental pipeline to identify and color code myeloid cells in the bone marrow and to generate random distributions. Scale bar = 200µm. **b.** Histograms showing the observed distribution of distances from each GMP (blue), MDP (green), MOP (orange) and GP (red) or random cells (white) to the closest indicated cell (n = 86 GMP from 4 sterna of 4 mice, n = 243 MDP from 23 sterna of 15 mice, n = 458 MOP from 11 sterna of 11 mice, n = 338 GP from 15 sterna of 12 mice). **c.** Representative image showing a Pre-neutrophil cluster lacking GP and the histograms showing the observed distribution of distances from each Pre-Neutrophil to the six closest Pre-Neutrophils (upper panel) or Immature Neutrophils (lower panels) in clusters with or without GP (n = 2493 Pre-Neutrophil from 3 sterna of 3 mice). Scale bar = 10µm. **d-i.** Histograms showing the observed (color) and random (white) distribution of distances from each Ly6C$^{lo}$ Mo (yellow), MDP (green), MOP (orange) and cDC (pink) to the six closest indicated cells d. n = 1603 Ly6C$^{lo}$ monocytes from 3 sterna of 3 mice; e. n = 67 MDP from 6 sterna of 4 mice; f. n = 171 MOP from 4 sterna of 4 mice; g. n = 1228 cDC from 6 sterna of 6 mice; h. n = 139 MDP from 11 sterna of 6 mice; **i.** n = 200 MOP from 5 sterna of 4 mice).

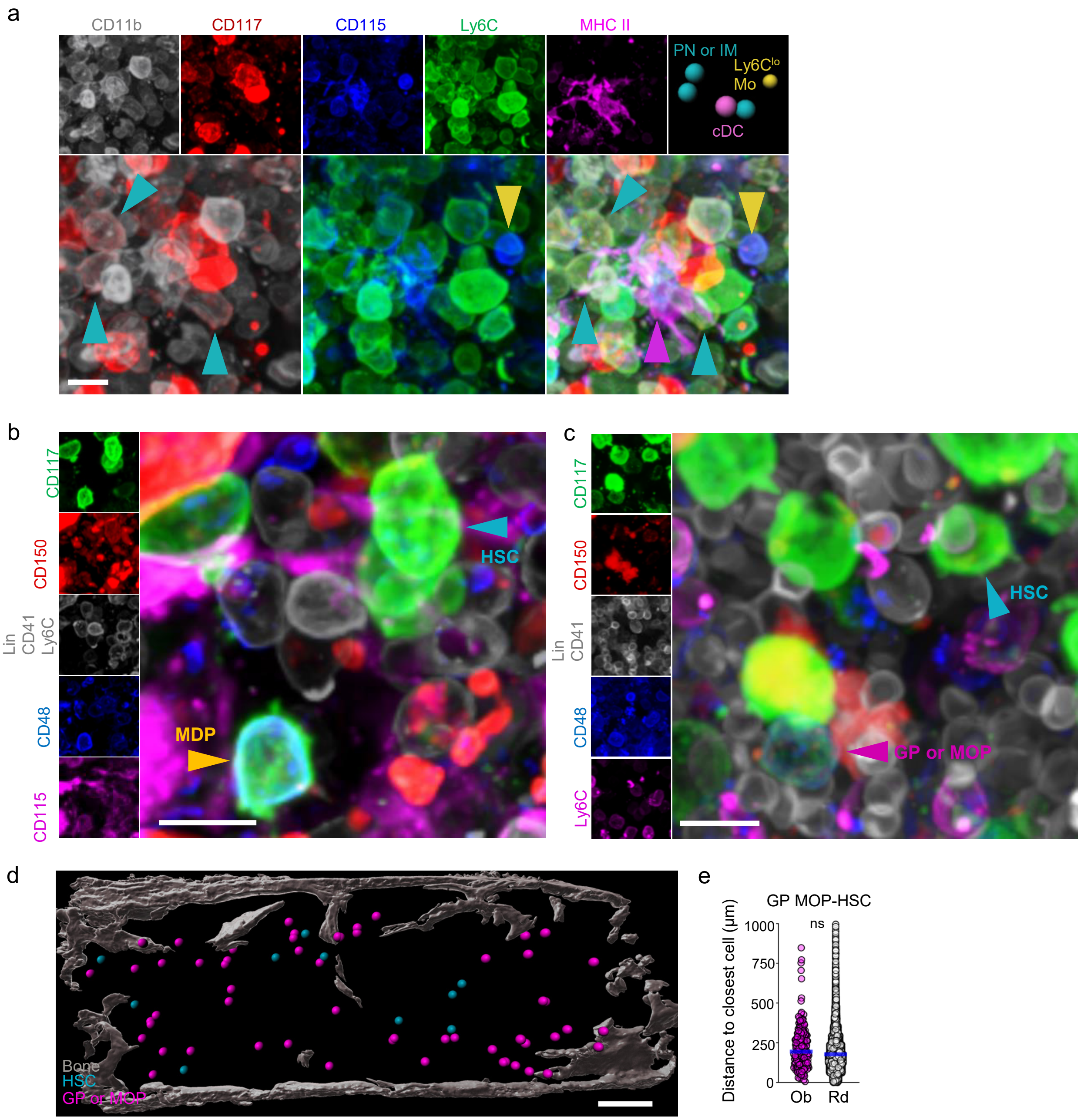


**Extended Data Figure 3. a.** Representative image showing simultaneous detection of Pre- and Immature Neutrophils, Ly6C$^{lo}$ monocytes and cDC. Scale bar = 10µm. **b.** Representative image showing detection of HSC and MDP in a single stain. Scale bar = 10µm. **c.** Representative image showing detection of HSC and a population containing CD117$^{+}$Ly6C$^{+}$ GP and MOP in a single stain. Scale bar = 10µm. **d.** Map showing the location of the HSC and GP/MOP cells in the bone marrow. **e.** Histograms showing the distance from each Lin$^{-}$CD117$^{+}$Ly6C$^{+}$ cells (either GP or MOP, pink dots) or the random simulation (white dots) to the closest HSC. Each dot corresponds to one cell and the dot radius is three times the average cell radius (n = 191 GP and MOP from 3 sterna of 3 mice).

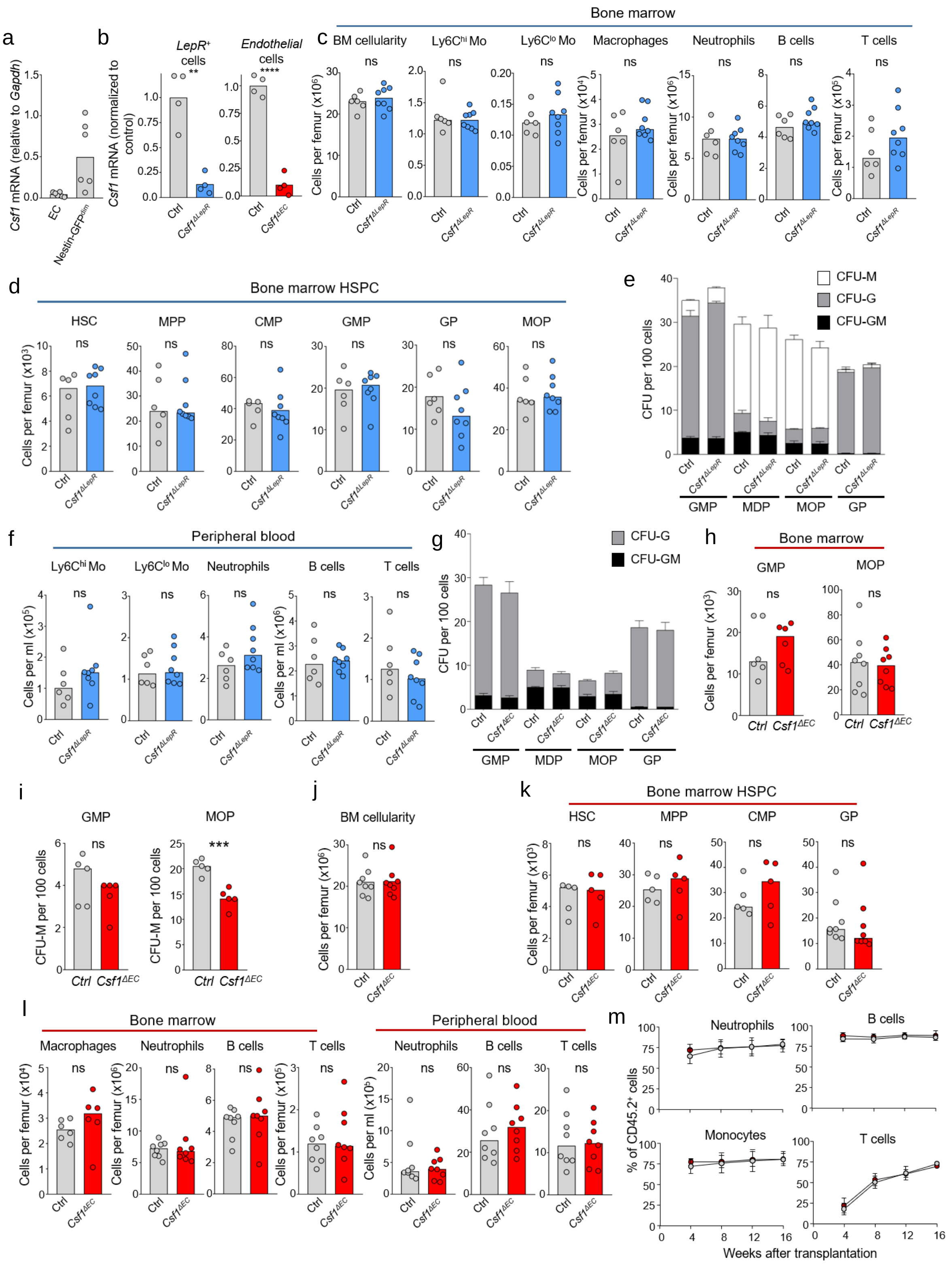

**Extended Data Figure 4. a.** qPCR analyses showing *Csf1* mRNA levels (relative to GAPDH) in FACS purified BM endothelial cells or Nestin-GFP$^{dim}$ perivascular cells (which largely overlap with LepR$^{+}$ perivascular cells[43]). **b.** qPCR analyses showing *Csf1* mRNA levels (normalized to Control) in FACS-purified CD45$^{-}$Ter119$^{-}$CD31$^{-}$LepR$^{+}$ cells or endothelial cells in control, *Csf1*$^{\Delta LepR}$, or *Csf1*$^{\Delta EC}$ mice. **c, d.** Number of BM cellularity or the indicated populations in the femur of control or *Csf1*$^{\Delta LepR}$ mice. **e.** Colony forming activity (d, black: CFU-GM, grey: CFU-G, white: CFU-M) of the indicated progenitors FACS-purified from control (n = 5) or *Csf1*$^{\Delta LepR}$ (n = 5) mice. **f.** Number of the indicated populations in the femur of control or *Csf1*$^{\Delta LepR}$ mice. **g.** Colony forming activity (black: CFU-GM, grey: CFU-G) of the indicated progenitors FACS-purified from control (n = 5) or *Csf1*$^{\Delta E}$ (n = 5) mice. **h, i.** Number (h) and CFU-M activity (i) of GMP or MOP from control or *Csf1*$^{\Delta EC}$ mice. **j-l.** Number of BM cellularity or the indicated populations in the femur or blood of control or *Csf1*$^{\Delta EC}$ mice**. m**. The panels show the percentage of the indicated CD45.2$^{+}$ cells in the blood of lethally irradiated CD45.1$^{+}$ recipients after transplant of $10^{6}$ BM cells purified from *Ctrl* ( white dots) or *Csf1*$^{\Delta EC}$ mice (red)- both CD45.2$^{+}$- together with $10^{6}$ CD45.1$^{+}$ competitor cells at the indicated time points after transplantation. The dots show the mean of 12 mice per group and the error bars show standard deviation. Unless otherwise indicated for all panels one dot corresponds to one mouse.

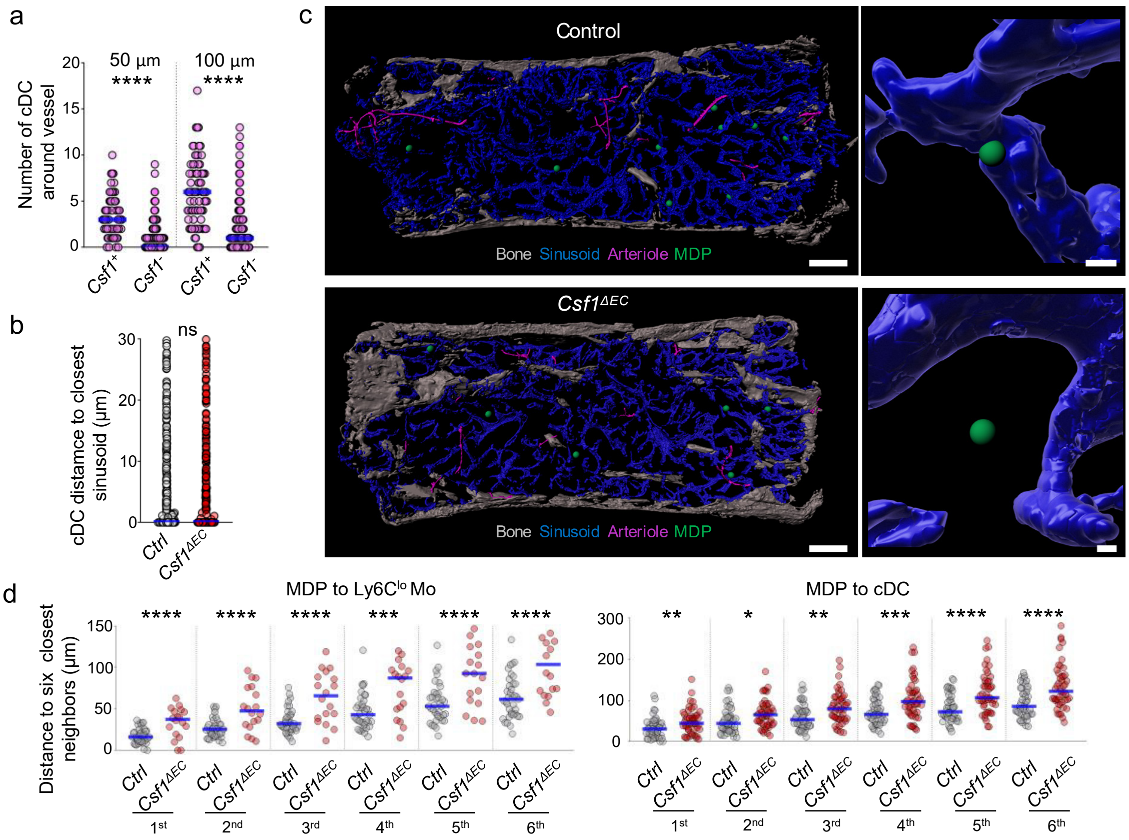


**Extended Data Figure 5. a.** Number of cDC found within the indicated distances of CSF1$^+$ and CSF1$^-$ vessels in wild-type (n = 76 CSF1$^+$ vessels and n = 520 CSF1$^-$ vessels in 4 sterna of 3 wild-type mice). **b.** Histograms showing the distance from each cDC to the closest sinusoid in control or $Csf1^{\Delta EC}$ mice (n= 451 cDC in 2 sterna of control mice, n = 343 cDC in 3 sterna of $Csf1^{\Delta EC}$ mice). **c.** Maps showing the relocation of MDP away from sinusoids in $Csf1^{\Delta EC}$ mice. Scale bars = 200 and 10µm. The radius of the dots is 3x (left map) or 1x (right images) the average radius of the MDP. **d.** Histograms showing the distribution of distances from each MDP to the six closest Ly6C$^{lo}$ Mo or cDC in control or $Csf1^{\Delta EC}$ mice (For MDP-Ly6C$^{lo}$ Mo, n = 37 MDP from 4 sterna of 3 Control mice, n = 18 MDP from 4 sterna of 3 $Csf1^{\Delta EC}$. For MDP-cDC, n = 47 MDP from 6 sterna of 3 Control mice, n = 47 MDP from 9 sterna of 3 $Csf1^{\Delta EC}$).
